# Effects of sensorimotor delays and muscle force capacity limits on the performance of feedforward and feedback control in animals of different sizes

**DOI:** 10.1101/2024.09.23.614404

**Authors:** Sayed Naseel Mohamed Thangal, Heather L. More, C. David Remy, J. Maxwell Donelan

## Abstract

Animals rely on both feedforward and feedback control for perturbation responses. When comparing animals of different sizes, we find that several features that affect perturbation responses change—larger animals have longer sensorimotor time delays, heavier body segments and proportionally weaker muscles. We used simple computational models to compare fast perturbation response times under feedforward and feedback control, as a function of animal size. We developed two tasks representing common perturbation response scenarios in animal locomotion: a distributed mass pendulum approximating swing limb repositioning (swing task), and an inverted pendulum approximating whole body posture recovery (posture task). First, we used a normalized feedback control system to show how feedback response times can either be limited by the force generation capacity of muscles (force-limited), or by sensorimotor delays which constrain the maximum feedback gains that can be used to produce stable responses (delay-limited). Next, we used more detailed scaled models which represent the full-size range of terrestrial mammals and parameterized the sensorimotor delays, maximum muscle forces, and inertial properties using published scaling relationships from literature. Across animal size and in both tasks, we found that feedback control was primarily delay-limited—the fastest responses used a fraction of the available muscle force capacity. Feedforward control, which is able to fully activate muscles and produce faster responses—was about four times faster than feedback control in the smallest animals, and around two times faster in the largest animals. For rapid perturbation responses, feedback control appears ineffective for terrestrial mammals of all sizes, as the fastest response times exceeded available movement times, while feedforward control did not. Thus, feedforward control is more effective for reacting quickly to sudden and large perturbations in animals of all sizes.

## 1. Introduction

Animals rely on both feedforward and feedback control strategies to respond to disturbances they encounter when moving through the world. Feedback control uses sensory input not only to sense the initial disturbance, but also to continuously compare the body’s movement to the desired movement throughout the response. The controller uses this changing error signal to generate the motor commands that continuously correct for the effect of the disturbance [1,2]. One benefit of this approach is that by using sensory feedback to continuously compare the desired movement to the actual movement, the disturbance can be accurately counteracted without advance knowledge of the disturbance’s effects. In contrast, feedforward control uses predefined motor commands to counteract the effect of a disturbance rather than continuously updating its response based on sensory feedback [3–5]. One benefit of this strategy is that it can be used to rapidly respond to a perturbation as it does not have to wait for continuous adjustment by sensory feedback. Animals use neural controllers that combine these two strategies, perhaps to benefit from their individual advantages [5–7].

In in-vivo studies of perturbation responses, the control strategy varied depending on the animal species and type of perturbation. Daley and colleagues conducted a comprehensive series of experiments to understand perturbation responses in running guinea fowl. The perturbations included potholes, step up, and step down obstacles; these perturbations were significant enough to force the bird to alter its gait, but allowed the bird to continue running and did not cause a fall. They found a consistent set of strategies that the birds used to navigate these perturbations. The birds maintained a constant swing leg angular cycling rate through rhythmic feedforward control of the proximal leg joints, irrespective of whether they were able to anticipate the perturbation or not. The distal joints then responded preflexively to compensate for the perturbation, acting either as a spring or a damper depending on the leg posture at heelstrike. Feedback modulation of the leg only occurred towards the very end of the perturbed stance phase [7–11]. Farley and colleagues studied human responses to anticipated or unanticipated changes in floor stiffness while hopping and running [12,13]. They showed that humans adjust the overall limb stiffness under feedforward control and rely on preflexes to dampen out perturbations. Similarly, studies on cockroaches have also shown that they rely on feedforward control for navigation [14,15]. On the other hand, Ting and colleagues studied postural responses to support surface translation and rotation perturbations, and showed that a time delayed feedback control model under Proportional-Derivative-Accelerative (PDA) control can reproduce the EMG signals of the postural response [16–18].

The performance of feedback and feedforward control systems is limited by time delays and actuator force capacity. The limited speed at which signals are conducted along nerves and across synapses results in sensorimotor time delays [19]. These time delays are not just present in biological control systems—electrical signal transmission and data processing time delays constrain the performance of engineering control systems [20]. Additionally, there is a limit to the maximum forces that animal muscles can produce [21], which parallel the force saturation limits seen in engineered actuators [20]. Time delays and actuator force capacity are two well studied limitations in engineering control systems [22–24]. Actuator force capacity can reduce the responsiveness of both feedforward and feedback control systems because the speed at which a physical system can be repositioned depends on how quickly the system can be accelerated, and this acceleration depends on the force capacity of its actuators. While time delays increase the time it takes to initiate a response in both feedforward and feedback systems, feedback systems have an additional vulnerability to time delays. When feedback control signals are outdated due to time delays, the motor commands generated are inappropriate for the current state of the system, resulting in poor control and reduced stability [1,25]. If the delays grow too large, the delayed feedback signal will actively destabilize the system instead of controlling it. To compensate for long feedback delays, a feedback-delayed controller must use lower gains to remain stable when compared to a controller with shorter time delays. This results in lower forces and slower responses. The main limiting factor in a feedback control system could either be its time delays or its actuator force capacity. A feedback system that doesn’t suffer from time delays will rapidly reach the force capacity of its actuators and how quickly it can respond will then depend upon how strong the actuators are at full capacity. In contrast, a feedback system that has long time delays, will never be able to tap into the full force capacity of its actuators because the long delays will necessitate low feedback gains to insure stability. Whether a feedback control system is force-limited or delay-limited will depend on the relative magnitudes of actuator strength and time delay duration.

We suspect that the effects of time delays and actuator force capacity on control performance depends on animal size. Sensorimotor delays are longer in larger animals when compared to smaller animals, even when expressed relative to movement time [19]. While larger animals have larger and stronger muscles when compared to smaller animals, their body segments are also heavier rendering their muscles proportionally weaker [19,26–29]. But working to the advantage of large animals is that the time available to make a corrective response gets longer—larger animals take longer to fall to the ground when they lose balance, and they have more time available to correct a disturbance within a stride when running [29]. Because these features scale with animal size, whether feedback control is limited by time delays or force capacity may depend on animal size. And the ability to control movement using feedforward and feedback control, as well as the relative advantage of one control approach over the other, may also depend on animal size.

Here our objective was to understand the performance of feedforward and feedback control strategies in animals of different sizes when constrained by sensorimotor delays and muscle force capacity limits. To accomplish this, we developed simple feedforward and feedback control systems, and scaled them to represent the size range of terrestrial mammals. The consequences of time delays and actuator force capacity are most evident when a system is required to respond as fast as possible, because fast responses require using the maximum possible muscle forces while ensuring that the response remains accurate and stable. Two fast response situations in animals are repositioning the swing leg after a stumble when running at high speed, and counteracting an aggressive push when standing still. Consequently, we quantified response times for two perturbation response tasks: a swing leg repositioning task (swing task), and a posture recovery task (posture task). We parameterized the models’ anatomical features based on animal scaling data from the literature and optimized their controller parameters to achieve the fastest responses. We simulated perturbations responses in these parameterized models to estimate the scaling relationship between response time and animal size for both feedforward and feedback control. By comparing response times under these two different types of control to each other, and to available movement times, we quantified the effectiveness of these control strategies across animal size and determined whether response times are limited by time delays or force capacity.

## 2. Methods and Models

To understand perturbation responses in animals, we developed two types of models with different levels of complexity (scaled models and a normalized feedback control model), considered two limitations to control performance (sensorimotor delays and muscle force capacity limits), and simulated two perturbation response scenarios (swing task and posture task). The scaled models are more detailed, consider gravity, simulate feedforward and feedback control, and are parametrized to represent the size range of terrestrial mammals. The normalized feedback model is a simplified version of the scaled feedback models that has only a single free parameter, allowing us to more clearly evaluate how sensorimotor delays and muscle force capacity limits interact to affect perturbation responses.

### 2.1 Scaled models

We developed scaled models that can represent the size range of terrestrial mammals using pendulums to represent the body segments, and parametrized them using data for inertial, muscular and neural features from the scaling literature. Fig 1 provides a schematic overview of the feedforward and feedback control system models.

**Fig 1.**
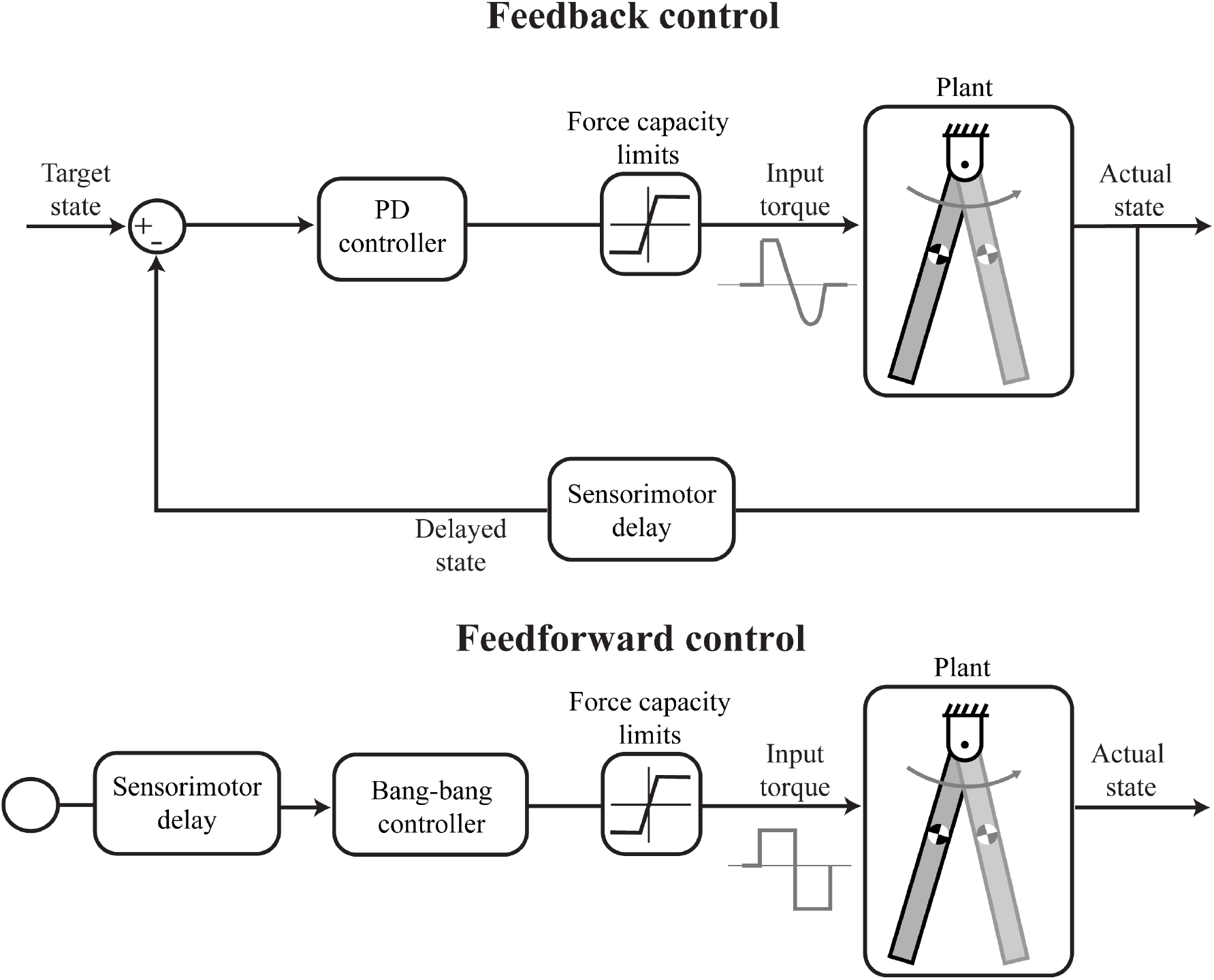
Feedback and feedforward control systems for the scaled models. The feedback system (top) uses a proportional-derivative (PD) controller to generate torque based on time delayed state feedback. The feedforward system (bottom) uses a bang-bang controller to generate torque. The plant represents the rigid body dynamics of the body segments using pendular models. The sensorimotor delay delays the sensory feedback about the plant state from reaching the controller, including the initial detection of the perturbation. The force capacity limits restrict the maximum positive and negative torques that can be applied by the controller, based on the force capacity of the muscles used in the task.

We made several assumptions to simplify the scaled models. For example, we assumed that the perturbation responses are mediated through monosynaptic spinal reflex pathways, and do not involve higher level (supraspinal or cortical) neural centers. While the term feedforward control is also used in the literature to refer to rhythmic signals generated by central pattern generators [7,30], we use feedforward control for predefined motor commands generated in response to a perturbation without using sensory feedback. We also simplified the scaled feedback models by assuming that all sensorimotor delays occur in the feedback (sensory) pathway. In real life, there are certain delays in the feedback (sensory pathway) and other delays in the feedforward (motor) pathway. In the supplementary materials section S1, we show how this assumption does not affect outcomes for the step response inputs we use in our simulations.

#### 2.1.1. Swing Task

##### Plant dynamics

The swing task represents an animal that encountered a trip of the forelimb during early swing phase, and has to reposition its foot by swinging it forward to avoid a fall; this is similar to the stumble corrective response in animals [31,32] and the elevating strategy in humans [33]. We modeled the forelimb as a distributed mass pendulum and the task required repositioning the limb under muscle torques applied at the shoulder, from rest at an initial negative angle to rest at a final positive angle (Fig 3a). The equations of motion are given by:

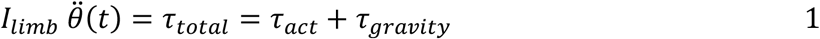

where *τ*_*act*_ is the torque exerted by the shoulder muscles, and *τ*_*gravity*_ the torque exerted by gravity. *I*_*limb*_ is the moment of inertia of the forelimb about the shoulder. 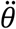 is the angular acceleration of the forelimb. We adopted the convention that the angle is zero when the limb is pointing vertically downwards and defined the counterclockwise direction to be positive. The torque due to gravity is described by:

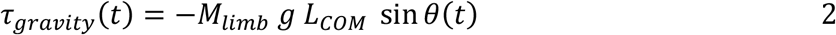

where M_l*imb*_ is the mass of the forelimb, and *L*_*COM*_ is the distance from the shoulder joint to the limb center of mass. *g* is the acceleration due to gravity (9.81 m/s^2^).

##### Feedback controller

Under feedback control, *τ*_*act*_ is the actual torque applied at the shoulder, and *τ*_*des*_ is the desired torque specified by the Proportional-Derivative (PD) controller output. *τ*_*act*_ and *τ*_*des*_ are not necessarily equal as *τ*_*des*_ is subject to force capacity limits:

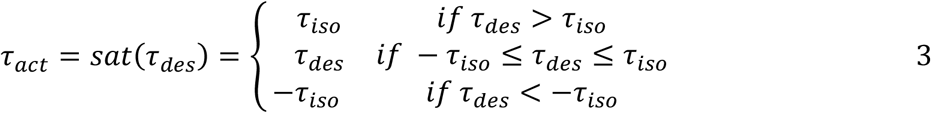

τ_*iso*_ is the force capacity limit, set as the maximum isometric torque that can be applied by muscles about the shoulder joint to reposition the swing limb. In this feedback controller, *τ*_*des*_ is defined as:

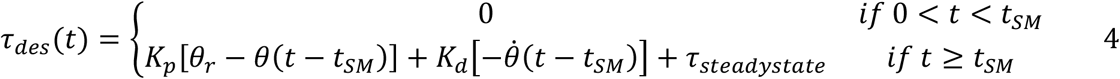

where *K*_*p*_ is the proportional gain, *K*_*d*_ is the derivative gain, *θ*_*g*_ is the reference angle (the final desired position of the limb), *τ*_*steadystate*_ is the steady state torque required to counter gravity at the final state, and *t*_*SM*_ is the sensorimotor time delay. For a duration equal to the sensorimotor delay at the beginning of the simulation, the controller does not apply any forces, and the forelimb moves under the influence of gravity. Once this deadtime is over, the controller begins to exert forces on the pendulum, using time-delayed state information.

##### Feedforward controller

Under feedforward control, the plant dynamics are identical but the applied torque (*τ*_*act*_) represents the torque produced by a bang-bang controller, with an initial deadtime:

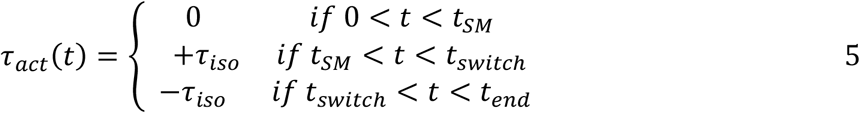

where *t*_*switch*_ is the time at which the controller changes the direction of applied torque. Table 1 summarizes the parameters used in this swing task scaled model, as well as how we determined the values.

**Table 1.** Table of input parameters, optimized parameters and results for the swing task. The Source column indicates whether the parameter was an input with values calculated from the scaling literature, or whether the parameter or performance measure was found through optimization. The Coefficient and Exponent columns are the coefficients and exponents in the power law equations that quantify the effect of size on that parameter.

| Parameter | Source | Coefficient (*a*) | Exponent (*b*) |
| --- | --- | --- | --- |
| **Swing task inputs** |  |  |  |
| $t_{SM}$: Sensorimotor delay (ms) | Literature | 31 | 0.21 |
| $M_{limb}$: Forelimb mass (kg) | Literature | $5.8 \times 10^{-2}$ | 1.00 |
| $I_{limb}$: Forelimb inertia (kg.m<sup>2</sup>) | Literature | $2.52 \times 10^{-4}$ | 1.75 |
| $L_{COM}$: COM length (m) | Literature | $5.6 \times 10^{-2}$ | 0.36 |
| $\tau_{iso}$: Triceps torque (N.m) | Literature | 0.54 | 1.19 |
| **Swing task outputs** |  |  |  |
| Swing duration at max sprint (ms) | Literature | 147.9 | 0.17 |
| **Feedback control results** |  |  |  |
| $K_p$: Proportional gain (N.m/rad) | Optimization | $1.75 \times 10^{-2}$ | 1.28 |
| $K_d$: Derivative gain (N.m/(rad/s)) | Optimization | $4.76 \times 10^{-3}$ | 1.54 |
| Feedback response time (ms) | Optimization | 198.9 | 0.21 |
| **Feedforward control results** |  |  |  |
| $t_{switch}$: Switch time (ms) | Optimization | 46.3 | 0.23 |
| Feedforward response time (ms) | Optimization | 61.9 | 0.24 |

##### Perturbations

In the feedforward control responses, the response time depends on the size of the movement—small movements are accomplished quickly, but large movements take more time. Perhaps non-intuitively, response time is independent of movement size in feedback control if the controller gains are the same and the torques do not exceed force capacity limits—it takes the same amount of time to perform a small movement as a large movement. Thus, to compare response times in the two types of control, it is useful to pick a movement size. Here we used a swing leg repositioning from -15° to +15° because sensorimotor delays and inertial delays are equally matched in a one kg animal for this movement size [27].

##### Simulations and optimization – feedback control

For the feedback control simulations, which required time-delayed state information, we used the dde23 delay-differential equation solver (explicit Runge-Kutta algorithm with discontinuity tracking) to numerically integrate the equations of motion (MATLAB R2020a, The MathWorks, Inc., Natick, MA, USA). We calculated both settling time and overshoot on the angle curve (Fig 3b). We quantified response time as the settling time with 2% thresholds at the reference angle (*θ*_*r*_) (Fig 3b). We constrained overshoot of the angle curve to be zero, to match a critically damped step response [34]. We normalized the simulation time by simulating for 20 *t*_*SM*_. To counter the steady state gravitational torques at the target position of +15°, we explored the use of a steady state torque (*τ*_*steadystate*_), or integral control. We used the steady state torque option because it required fewer parameters to optimize and produced faster response times. We used MATLAB optimization functions, fminsearch (simplex algorithm) and fmincon (interior-point algorithm) to search for the ideal parameters for the feedback controller (*K*_*p*_, *K*_*d*_) that produced the fastest response times without overshoot.

##### Simulations and optimization – feedforward control

As the feedforward simulations did not require time-delayed state information, we used MATLAB’s ode45 variable time step solver (explicit Runge-Kutta algorithm) to numerically integrate the equations of motion. Under feedforward control, we used event detection to terminate the simulation when the pendulum came to rest and used the elapsed time as response time. We used optimization to find the ideal parameter for the feedforward controller (*t*_*switch*_) that stopped the forelimb at the reference target, while achieving the fastest response time.

##### Scaling relationships

Two recent publications that quantified the scaling of mammalian anatomical and neural features, and an older publication on the scaling of muscle features in mammals, provide us with sufficient data to parametrize our scaled control systems [19,26,28]. We used data from More and Donelan (2018) on the monosynaptic stretch reflex pathways to parametrize the sensorimotor delay(*t*_*SM*_) [19]. We used data from Kilbourne and Hoffman (2013) on terrestrial mammalian limbs to set inertial properties in this model (*M*_*limb*_, *L*_*COM*_, *I*_*limb*_) [28]. We estimated the maximum force capacity of the shoulder muscles (*τ*_*iso*_) using muscle mass, length and moment arm data for the triceps published by Alexander et al. (1981) [26]. We found muscle volume from mass by assuming a density of 1060 kg/m^3^ [35], and cross-sectional area by dividing muscle volume by muscle length. We did not consider muscle pennation, and its effects on cross-sectional area in our models. We then multiplied cross-sectional area by the isometric stress of muscles, estimated to be 20 N/cm^2^ [21,36], to get muscle force. *τ*_*iso*_ is muscle force multiplied by the moment arm. We evaluated the models for eight logarithmically spaced animal sizes from 1 gram to 10 tons to cover the size range of terrestrial mammals, and also included 5 grams and 5 ton sizes for 10 total animal sizes [37,38]. We then performed a least squares linear regression on the logarithmically transformed animal mass vs. simulation output data (controller gains and response times) to extract the coefficient and exponent of the underlying scaling relationship [39].

#### 2.1.2 Posture task

##### Plant dynamics

The posture task represents an animal recovering its posture after a push forward in the sagittal plane, under the control of its plantarflexors [16,17,40–42]. We used a point mass inverted pendulum to represent the entire body, with a mass *M* equal to the weight of the whole animal, and length *L* set to the leg length for each animal mass (Fig 5a). We modeled forward push perturbations as impulsive forces that result in changes to the body’s initial forward velocity, but not it’s position. The task required generating torques to counter the initial velocity caused by the push and returning to rest at the vertical position from which the system started. The equations of motion are given by:

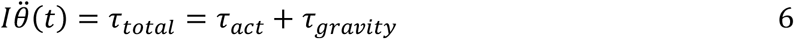

where *τ*_*act*_ is the torque exerted by the muscles, and *τ*_*gravity*_ the torque exerted by gravity. *I* is the moment of inertia of the pendulum, computed as *ML*^2^. In contrast to the swing task, we defined the angle to be zero when the inverted pendulum is pointing vertically *upwards*, and defined the counterclockwise direction to be positive. The torque due to gravity is consequently defined as:

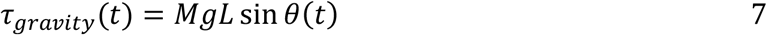

The feedback controller in the posture task did not require a steady state term, as there are no gravitational torques at the vertical end position. Other than this difference, we determined *τ*_*act*_ for feedforward and feedback control in the posture task using identical equations to the swing task (Eqns 3,4,5). Table 2 summarizes the parameters used in this posture task scaled models, as well as how we determined the values.

**Table 2.**
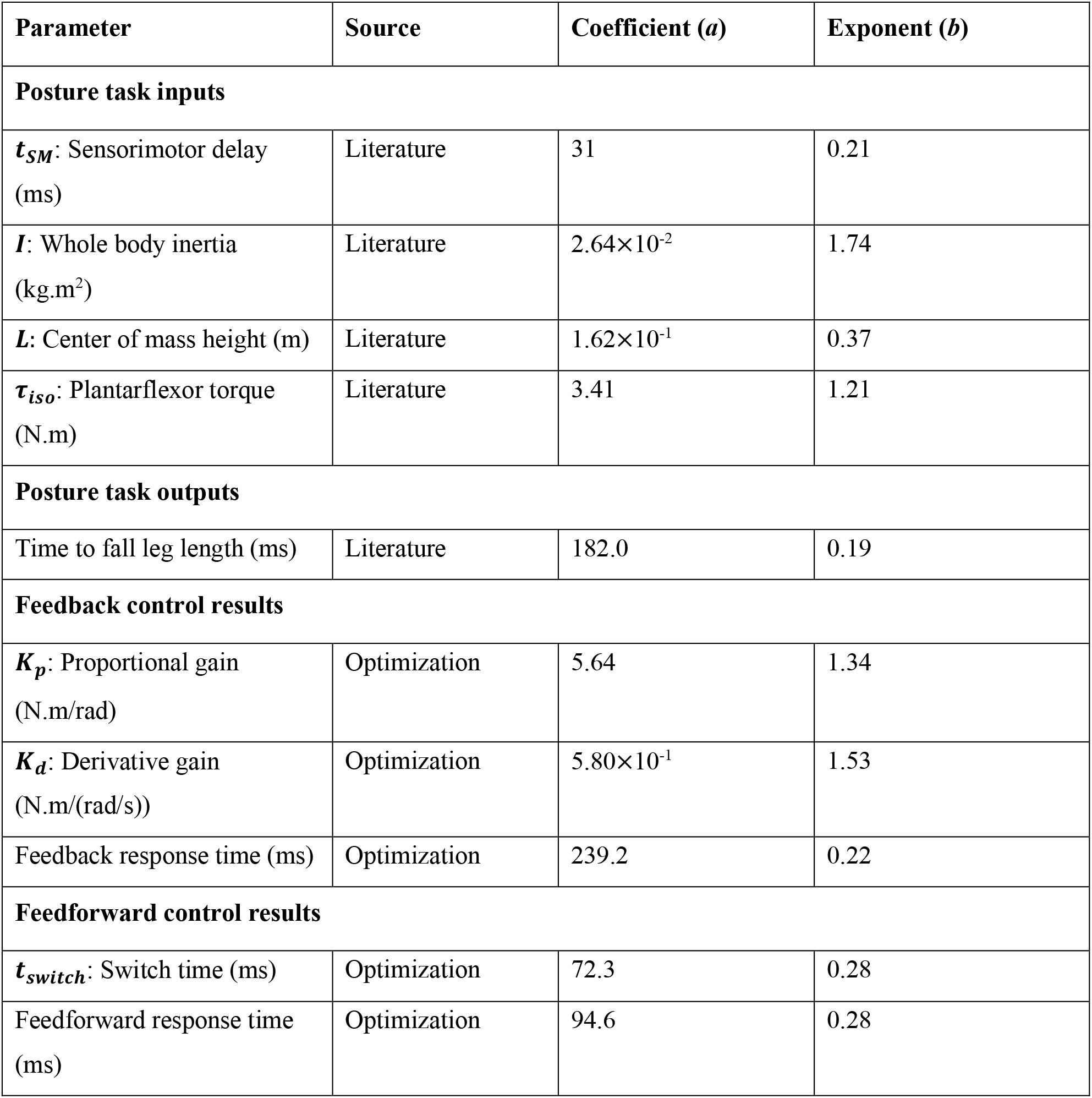
Table of input parameters, optimized parameters and results for the posture task.

| Parameter | Source | Coefficient ($a$) | Exponent ($b$) |
| --- | --- | --- | --- |
| **Posture task inputs** |  |  |  |
| $t_{SM}$: Sensorimotor delay (ms) | Literature | 31 | 0.21 |
| $I$: Whole body inertia (kg.m<sup>2</sup>) | Literature | $2.64 \times 10^{-2}$ | 1.74 |
| $L$: Center of mass height (m) | Literature | $1.62 \times 10^{-1}$ | 0.37 |
| $\tau_{iso}$: Plantarflexor torque (N.m) | Literature | 3.41 | 1.21 |
| **Posture task outputs** |  |  |  |
| Time to fall leg length (ms) | Literature | 182.0 | 0.19 |
| **Feedback control results** |  |  |  |
| $K_p$: Proportional gain (N.m/rad) | Optimization | 5.64 | 1.34 |
| $K_d$: Derivative gain (N.m/(rad/s)) | Optimization | $5.80 \times 10^{-1}$ | 1.53 |
| Feedback response time (ms) | Optimization | 239.2 | 0.22 |
| **Feedforward control results** |  |  |  |
| $t_{switch}$: Switch time (ms) | Optimization | 72.3 | 0.28 |
| Feedforward response time (ms) | Optimization | 94.6 | 0.28 |

##### Perturbations

As in the swing task, we had to choose a perturbation size to compare feedback and feedforward response times. To trigger postural perturbations of similar magnitudes in different sized animals, we scaled the initial linear velocity for the inverted pendulum mass with a dimensionless velocity of 0.21 Froude number [43]. At this perturbation size, the sensorimotor delays equaled the inertial delays in a one kg animal [27]. We defined the target angle and velocity to be zero, to bring the pendulum to rest at the vertical position.

##### Simulations and optimization-feedback control

Under feedback control, as the posture task involved a velocity perturbation, we calculated settling time with 2% thresholds on the angular velocity curve (Fig 5b). As overshooting the vertical position during a posture correction can also result in loss of balance, we calculated overshoot on the angle curve and constrained it to be zero during the optimization.

##### Simulations and optimization-feedforward control

Under feedforward control, we used event detection to terminate the simulation when the inverted pendulum came to rest, and optimized *t*_*switch*_ to stop the pendulum at the vertical position. Apart from these differences, we simulated and optimized the simulations using identical methods to the swing task.

##### Scaling relationships

We used the average length of the mammalian forelimb and hindlimb based on data from Kilbourne and Hoffman (2013) to set inverted pendulum length [28]. We set *τ*_*iso*_ to equal four times the maximum torque that can be exerted by the ankle plantarflexors based on Alexander et al. (1981) [26].

### 2.2 Normalized feedback control system

In addition to the family of scaled feedback models we describe above, we developed a single normalized model to study how feedback control can be limited by time delay and force capacity. This normalized feedback model is a simplified version of the scaled feedback models and only has a single free parameter, allowing us to more clearly evaluate how sensorimotor delays and muscle force capacity limits interact to affect perturbation responses. This single model applies to both swing and posture tasks in animals of all sizes. To accomplish this, we simplified the scaled feedback models by ignoring gravity, thereby reducing the plant to a double integrator under feedback control. Before normalization, the desired torque *τ*_*des*_ produced by the feedback controller is given by:

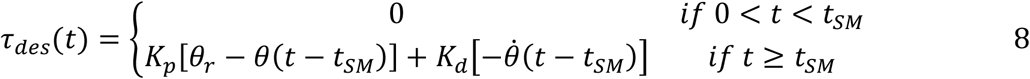

To non-dimensionalize Eqn 8 (same as Eqn 4 without the steady state torque component), we used three characteristic parameters of the perturbation response: *I* is the moment of inertia of the body segments being repositioned, *θ*_*r*_ is the reference target for feedback control and represents the size of the perturbation in the swing task, and *t*_*SM*_ is the sensorimotor time delay. We divided each torque component in Eqn 8 by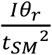, where *I* has fundamental units of M^1^L^2^, *t* _*SM*_ has units of T, and *θ*_*r*_ is dimensionless:

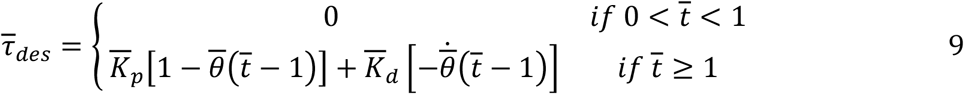

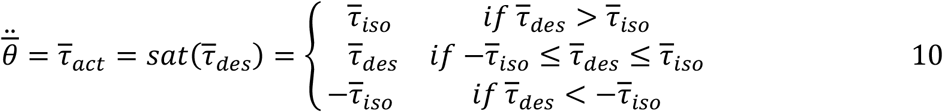

Eqns 9 and 10 represent the normalized feedback system, with all parameters and system behaviour expressed in dimensionless units. For example, the normalized proportional controller gain parameter is given by 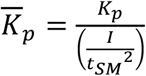 and the normalized response time result is given by 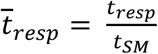. The force capacity, represented by 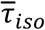, is the only free parameter in this model. Unlike in the scaled models, parameters related to animal size, time delay, and perturbation magnitude are all unity in this model. In all our models, including this one, controller gains are not free parameters as they are fixed at their optimal values (which we determined by optimization).

This model applies to both swing and posture tasks. We focus on studying the swing task in the main body of this manuscript and the posture task in the supplementary material. We also present additional analyses using Bode plots in the supplementary material that explains how time delays limit controller gains in a linear feedback system.

We simulated and optimized the normalized swing and posture task models using the same protocols and methods as described for the scaled models, except for the following differences.

For the swing task, we set the initial conditions for plant state 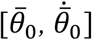 to [0,0], and the target state 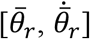 to [1,0]. We first simulated the normalized model without a force capacity limit 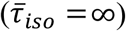, and then for a range of force capacity limits 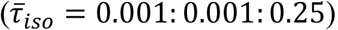, and optimized the controller gains (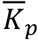 and 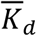) for each force capacity limit to achieve the fastest response time. We performed the derivations to normalize the equations by hand, and simulated the normalized feedback system using MATLAB’s Simulink toolbox.

## 3. Results

### 3.1 Normalized feedback control system with time delays and force capacity limits

Optimizing without force capacity limits 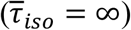 and with a dimensionless time delay of unity, we found that controller gains of 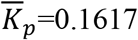 and 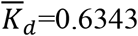 produced the fastest response time of 7.09, all in normalized units. The magnitude of this response time means that it took ∼7x the duration of the sensorimotor time delay to move the physical system from its original state to settle on its new target position. The torque during this simulation reached a maximum value of 0.1617, equal to the value of 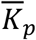 This maximum simulated torque means that while there was no limit to available force, the control system was only able to use 0.1617 units of dimensionless torque at a maximum. The optimal gains and system performance would be identical with a force capacity of 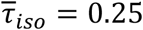, for example, as it would be without any limit to force capacity. Thus, we next systematically decreased 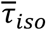 from 0.25 to 0, keeping time delay constant at unity, and optimized controller gains to achieve the fastest response times (Fig 2).

**Fig 2.**
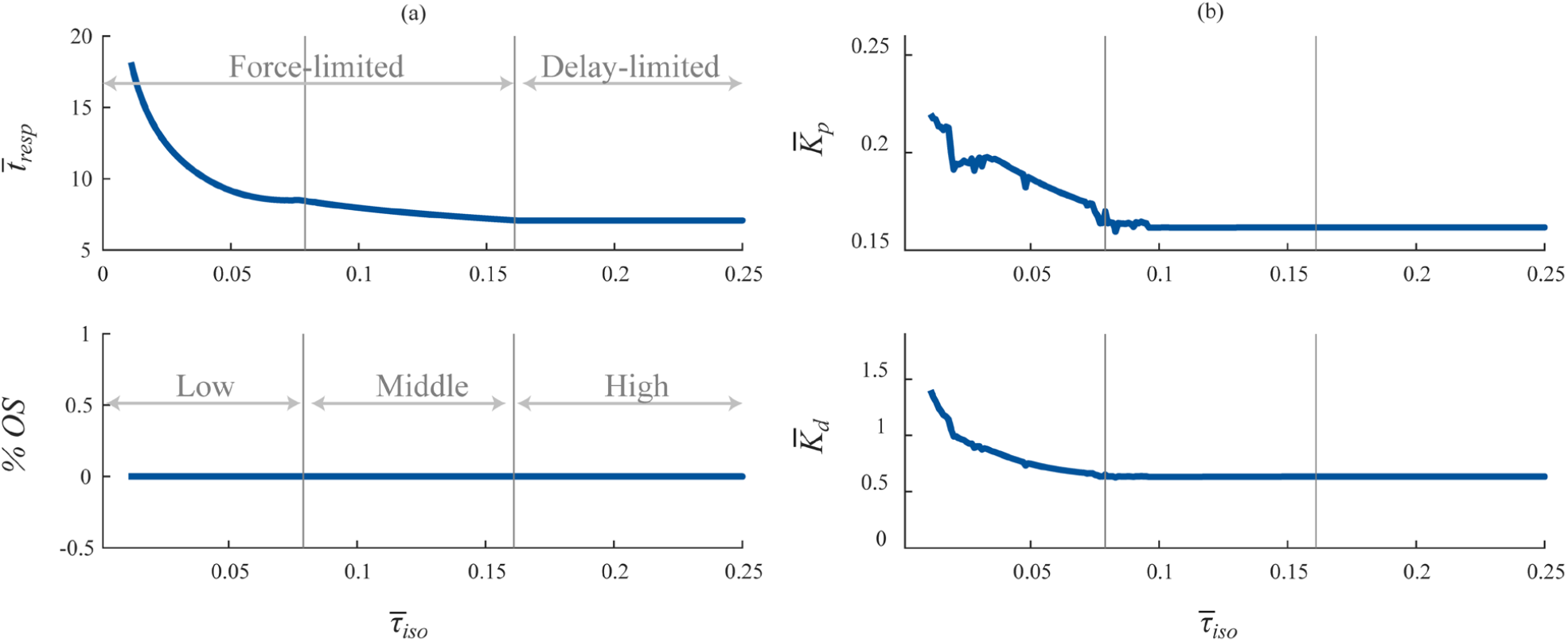
Normalized model—swing task response time vs. force capacity limits. (a) Changes in normalized response time 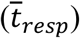 and overshoot (%OS) when force capacity limits 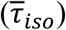 are lowered from 0.25 to 0, and time delay is kept constant at 1. There are three distinct regions in the response time curve indicated by the grey vertical lines: a high region 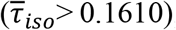, a middle region 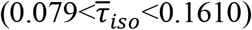, and a low region 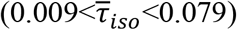. In the high region, feedback response time is purely delay-limited; the peak torques do not reach the force capacity limits. In the middle region, positive torques are clipped by the force limits, but not negative torques. In the low region, both positive and negative torques are clipped by the force limits. Thus, feedback response time is force-limited in the middle and low regions. (b) Changes in the optimal controller gains that produce the fastest responses for a range of force capacity limits. Note that there is some variability in the optimal controller gains in the middle and low regions due to multiple local minima that have values very close to the global minimum, but they do not affect the response time significantly.

This analysis shows that for feedback control systems with both time delays and actuator force capacity, there are two operating ranges: a delay-limited range and a force-limited range. Consider first the extremes. Without force capacity limits or delays, infinitely high gains can produce instant response times. However, if either the force capacity was set to 0, or if time delays were infinitely long, response times would be infinitely long. The delay-limited range consists of the high force capacity region 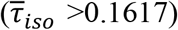, where only time delays affect response time. The force-limited range consists of the middle and low force capacity regions 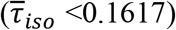, where force capacity limits begin to also limit response time. Extrapolating to animals, this analysis indicates that for an animal to respond quickly, it requires both strong muscles and short sensorimotor delays. Deficiencies in either factor can slow the animal’s ability to respond quickly. Whether an animal is delay-limited or force-limited would depend on the relative magnitudes of factors that affect the perturbation response—such as the moment of inertia of the body segments being moved, the sensorimotor delays, the size of the perturbation, and the muscle force capacity. These effects are seen in the normalization factor 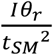 where increases in inertia or perturbation size will cause a shift towards the force-limited range even if the absolute force capacity is unchanged. And an increase in *t*_*SM*_, which is in the denominator of this factor, will cause a shift away from the force-limited range and towards the delay-limited range, again even if muscles have no increase in absolute force capacity. We found similar results for the posture task (supplementary materials section S3). We also compared the predictions from the normalized model to the results from the scaled model simulations and describe the results in section S4.

### 3.2 Scaling of perturbation response times for scaled models

#### 3.2.1 Swing task

Feedback control performed poorly when compared to feedforward control in the swing task (Figs 3 and 4). For a movement size of 30°, the fastest response times under feedback control scaled as 199*M*^0.21^ ms, compared to 62*M*^0.24^ ms under feedforward control (Fig 3c). This is because the feedback controllers were delay-limited, and unable to utilize a significant portion of their muscle force capacity due to the stability limitations imposed by long sensorimotor delays (Fig 4a). That the models were delay-limited across animal size means that the sensorimotor delay scaling determines the feedback control response time scaling—the feedback control scaling exponent is 0.21 because the sensorimotor delay exponent is 0.21. Because the exponents are the same, the feedback control response times are a constant multiple of the sensorimotor time delay and determined by the ratio of scaling coefficients (198.9/31). This equates to response time being 6.4x the duration of the time delay, which is close to the 7x prediction of the much simpler normalized model above. The feedforward control response is a combination of an initial deadtime during which the limb falls under gravity, and a bang-bang controlled repositioning movement under maximal muscle torques (*τ*_*iso*_). While the initial deadtimes scaled with sensorimotor delay (*M*^0.21^), the bang-bang movement scaled approximately with inertial delay (*M*^0.28^) [27]—together causing feedforward response times to scale with *M*^0.24^. Therefore, the ratio of feedback response times to feedforward response times gets smaller with animal size, ranging from four in smaller animals to two in larger animals (Fig 4b). Table 1 lists the input parameters and the optimized output parameters in the swing task.

**Fig 3.**
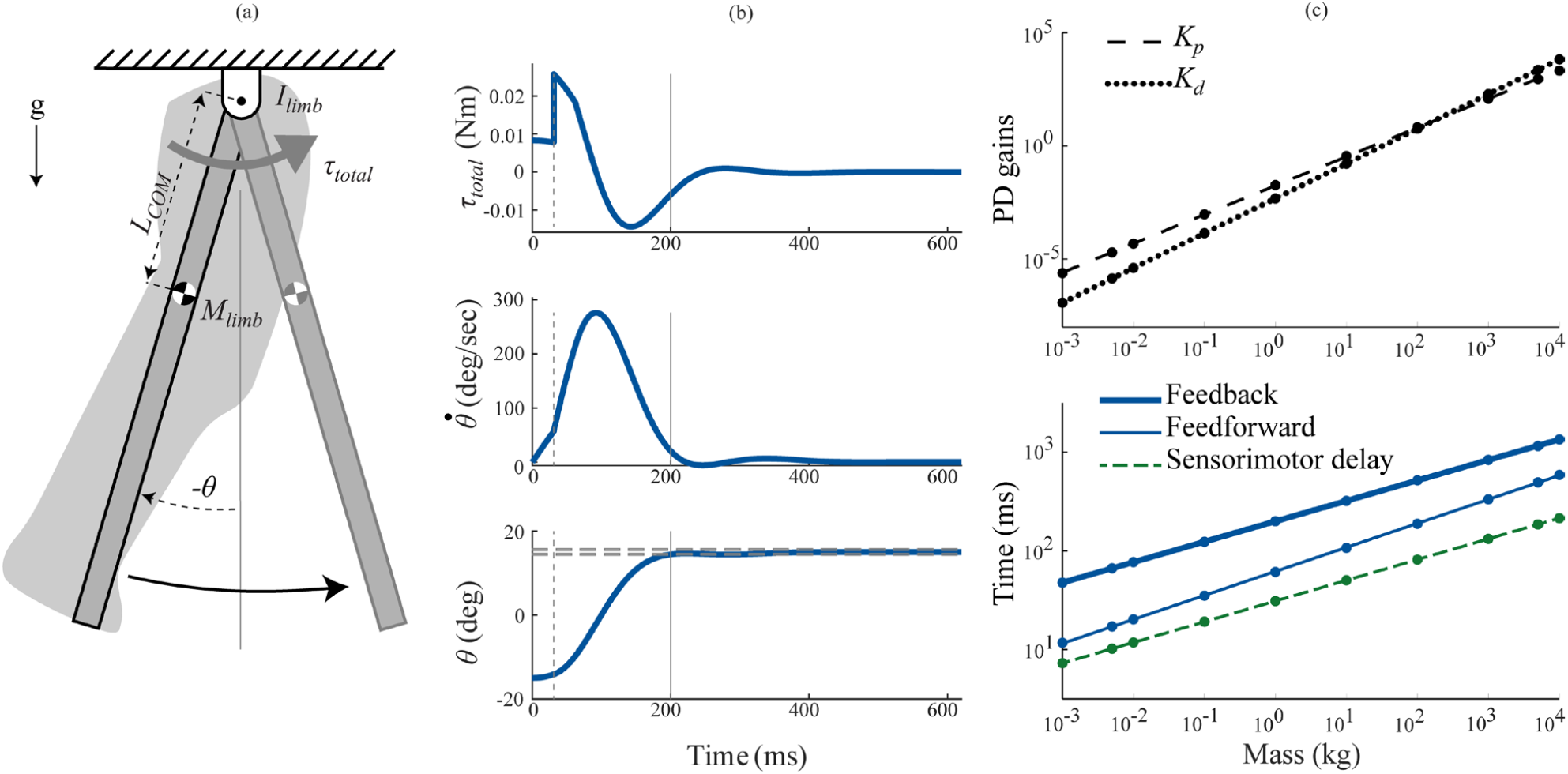
Scaled model—swing task responses under feedback control. The swing task simulates repositioning of the forelimb in response to an early swing phase trip. (a) We modeled the plant as a distributed mass pendulum actuated by torques generated by a PD controller. (b) Torque (*τ*_*total*_), angular velocity 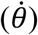 and angle (*θ*) profiles in the swing task for a one kg animal and a 30° movement under feedback control. The grey dashed line at 31 ms represents the initial sensorimotor delay period and the gray vertical line at 200 ms represents response time, computed as the settling time of the angle curve with a 2% threshold (grey dashed horizontal lines). (c) Log-log plots for the scaling of the controller gains (top) and the response times under feedback control (thick blue line), feedforward control (thin blue line) and sensorimotor delays (thin dashed green line). The dots denote the actual values obtained through optimization, while the lines denote the power law fit.

**Fig 4.**
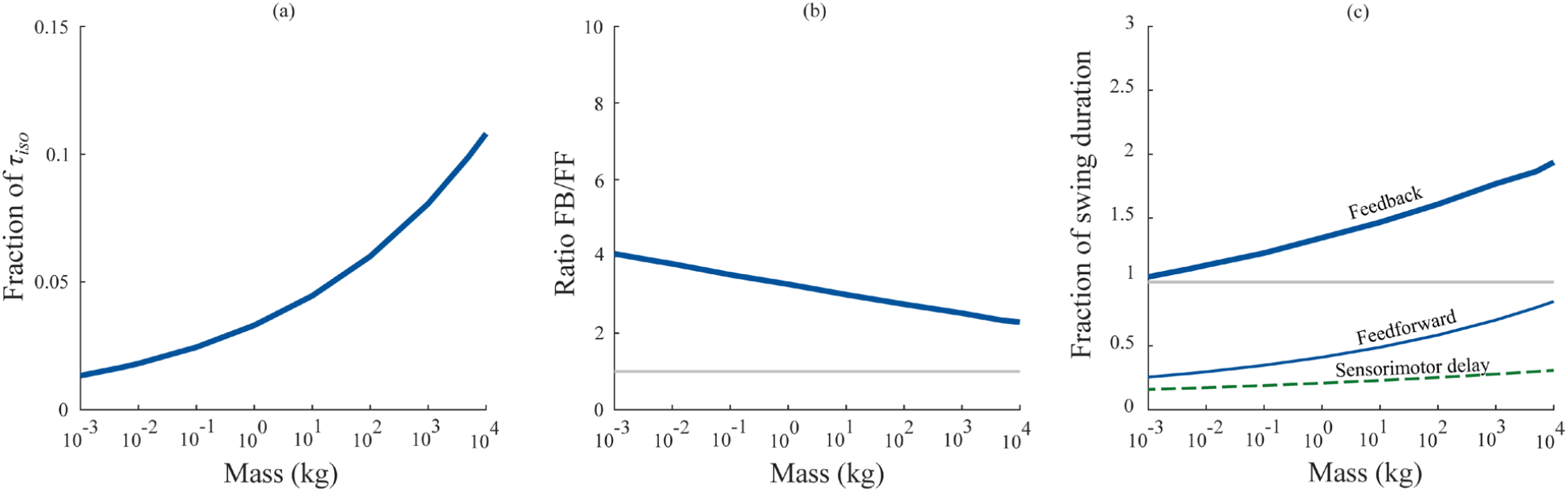
Swing task—comparison of feedforward and feedback response times. (a) The fraction of torque available based on muscle force capacity (*τ*_*iso*_) used by the feedback controller in the swing task across animal size. (b) The ratio of feedback control response times to feedforward control response times. (c) Fraction of swing duration at maximum sprint speed required to perform a corrective movement using feedback control (thick blue line), feedforward control (thin blue line) and sensorimotor delays (thin dashed green line).

Larger animals also have slower characteristic movements, giving them longer to complete a perturbation response. We estimated the shortest time available to complete a perturbation response for animals of different sizes, as the swing duration at maximum sprint speed [19,27]. Response times under feedback control exceeded swing duration at maximum sprint speed for all animal sizes, while feedforward control did not (Fig 4c).

#### 3.2.2 Posture task

While the feedback controllers in the posture task were able to use more of the available torque than in the swing task, it still performed poorly compared to feedforward control (Figs 5 and 6). For a 0.21 dimensionless velocity perturbation, the fastest response times under feedback control scaled as 239*M*^0.22^ ms, compared to 95*M*^0.28^ ms under feedforward control (Fig 5c). Other than for the largest animals, feedback control in the posture task was delay-limited rather than force-limited (Fig 6a). The feedback control response time was approximately 7.7x the duration of the sensorimotor time delay across animal size. Under feedforward control, initial deadtime scaled with sensorimotor delay (*M*^0.21^), and the bang-bang repositioning movement scaled with inertial delay (*M*^0.35^) [27]—together causing feedforward response times to scale with *M*^0.28^. The ratio of feedback response times to feedforward response times ranged from four in smaller animals to one and a half in larger animals (Fig 6b). Table 2 lists the input parameters and the optimized output parameters in the posture task.

**Fig 5.**
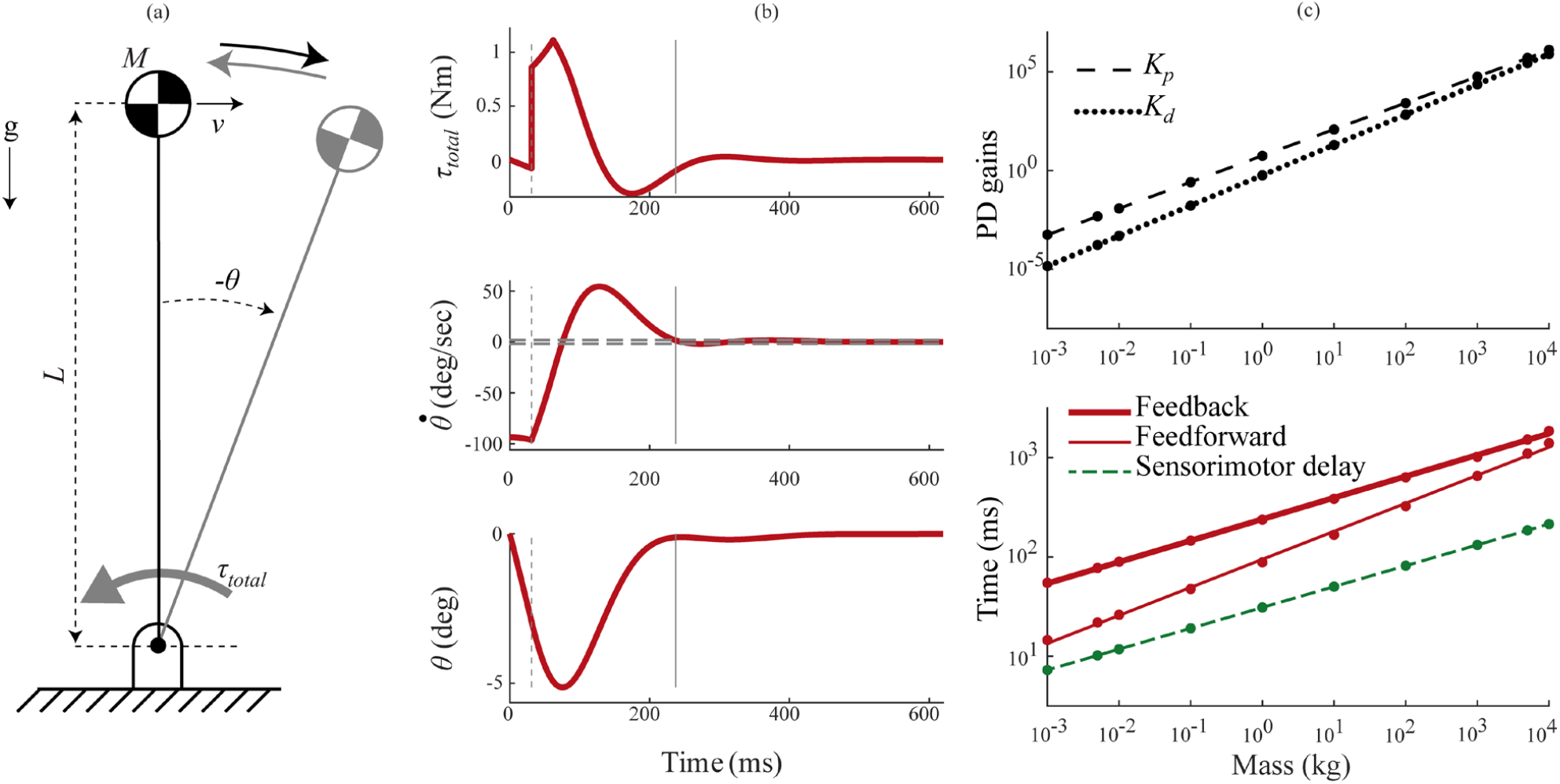
Scaled model—posture task responses under feedback control. The posture task simulates whole body posture control after a push forward in the sagittal plane. (a) We modeled the plant as a point mass pendulum actuated by torques generated by a PD controller. (b) Torque (*τ*_*total*_), angular velocity 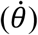 and angle (*θ*) profiles in the posture task for a one kg animal, perturbed by a force causing an initial dimensionless velocity of 0.21, under feedback control. The grey dashed line at 31 ms represents the initial sensorimotor delay period and the gray vertical line at 239 ms represents response time, computed as the settling time of the angular velocity curve with 2% thresholds (grey horizontal dashed lines). (c) Log-log plots for the scaling of the controller gains (top) and the response times under feedback control (thick red line), feedforward control (thin red line) and sensorimotor delays (thin dashed green line). The dots denote the actual values obtained through optimization, while the lines denote the power law fit.

**Fig 6.**
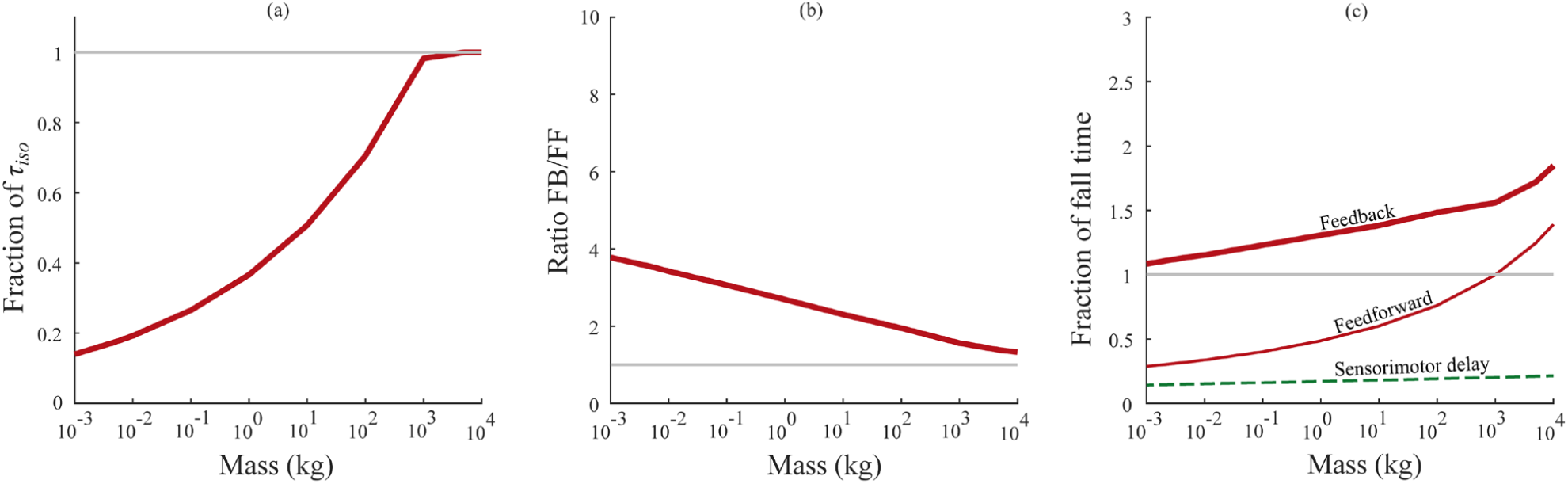
Posture task—comparison of feedforward and feedback response times. (a) The fraction of maximum isometric torque (*τ*_*iso*_) utilized by the feedback controller in the posture task across animal size. (b) The ratio of feedback control response times to feedforward control response times. (c) Fraction of available time (time taken to fall to the ground under gravity) required to perform a corrective movement using feedback control (thick red line), feedforward control (thin red line) and sensorimotor delays (thin dashed green line).

Feedback control response times in the posture task for the 0.21 dimensionless velocity perturbation also exceeded available time for all animal sizes. We estimated time available to complete the perturbation response as the time required to fall under gravity to the ground, from a height equal to the leg length for each animal size, which scaled as 182*M*^0.19^ ms. Feedforward response times for the largest animals also exceeded available time (Fig 6c). Feedforward response times get longer with perturbation size, while feedback response times do not (unless muscle force capacity limits are exceeded). If we had used stronger perturbations, we would generally have seen longer feedforward response times while feedback response times remained constant, resulting in smaller feedback to feedforward response time ratios (Fig 6b), and lighter animals exceeding available movement time.

We have included some additional analyses in the supplementary materials. Section S5 provides a breakdown of the components of the overall torque applied by the feedback controller in the scaled model simulations. Section S6 compares the scaled model results to in-vivo perturbation responses for cats and humans reported in the literature.

## 4. Discussion

In this study, we developed simple computational models to represent fast perturbation responses in animals, which are subject to sensorimotor delays and muscle force capacity limits. We quantified how response times scaled with animal size under feedforward and feedback control. Previous publications from our group had quantified the scaling of sensorimotor delays (signal transmission and processing time delays in the monosynaptic stretch reflex) [19], and inertial delays (movement time required to physically reposition body segments as part of the response to a perturbation) [27]. In this paper, we evaluated how the type of control used—feedforward vs. feedback control—further affected response times. We found that while feedforward control can fully activate muscles and produce fast responses, long sensorimotor delays required feedback control to use low gains to ensure stability, allowing only a fraction of the muscles’ force capacity to be utilized. That is, the effectiveness of feedback control within the size range of terrestrial mammals is predominantly delay-limited rather than force-limited. Feedback response times were about seven times longer than the duration of sensorimotor delays, and at least one and a half times longer than feedforward response times across animal size, for both the swing task and posture task. Feedback response times exceeded available movement time for all animal sizes, while feedforward response times did so only for the largest animals in the posture task (Figs 4c and 6c). Feedback control does not seem effective for fast perturbation responses in animals of any size.

To reach these conclusions, we had to make several assumptions and simplifications. We have carried over assumptions made in the estimation of parameters in previous publications, such as for the scaling of sensorimotor delays [19] and inertial delays [27]. For example, for the scaling of sensorimotor delay, More and Donelan (2018) had assumed that sensing delay, synaptic delay and neuromuscular junction delay are constant across animal size based on limited information in the literature. In calculating muscle force capacity limits, we assumed that the triceps (swing task) and the ankle extensors (posture task) are the dominant muscles involved in moving their respective joints, that their antagonistic muscles scale similarly, and that the isometric stress produced by mammalian muscle is constant at 20 N/cm^2^ [36]. To keep our models simple, and due to lack of data to accurately scale the muscle-tendon architecture across the size range of terrestrial mammals, we did not incorporate dynamic muscle models (e.g. Hill type). Although animal limbs are actually multi-jointed and multi-muscled, we used pendular models to represent body segments, and a single pair of opposing muscles to actuate them [26,44]. We have also not considered several animal features that have been shown to change with animal size such as posture, limb stiffness, joint damping and sensor accuracy; evaluating the effects of these features would require more complete neuromusculoskeletal models [45–47]. We used a proportional-derivative feedback controller as it is simple to implement, simple to interpret, and has been successfully used in the literature to model control of locomotion [16,48–51]. The actual control strategies implemented within the spinal reflex pathways is undoubtedly more complex, involving several control pathways with varying time delays, and remains a topic of research [52–55]. In engineering control systems, there are complex feedback controllers such as Model Predictive Control, or Smith predictors, which can compensate for time delays and generate bang-bang control. Here, we assumed that the spinal-level synapses that represent the feedback controller do not encode these complex algorithms and use the simpler and more straightforward proportional-derivative control instead.

Given these assumptions, as well as the nature of any purely modeling approach, these results are estimates that need to be tested experimentally. We have tried to estimate the limits of neural control by considering sensorimotor delays from the longest reflex pathways (a distance equal to twice the animal’s leg length), which use the least neural computation (a single synapse), and compared response times to the shortest available movement times (e.g. swing duration at maximum sprint speed). Using these highly simplified models to evaluate the limits of performance as a function of animal size, we have made predictions about the complex processes of animal locomotor control. One candidate experimental approach to test our predictions would be in-vivo experiments where the delays in sensory feedback or transmission of motor commands can be systematically manipulated while studying the response of the nervous system [56]. Several studies have investigated the effects of removing sensory feedback on locomotor behavior in animal models [7,31,49,57]. Instead of removing sensory feedback, one might decrease the nerve conduction velocity using cooling techniques to increase the sensorimotor delays in the reflex [58,59]. Such experiments would have to devise ways to cool only the nerves while minimizing changes to the muscle, perhaps by using nerve cuffs [60]. Experiments of this nature would be challenging to carry out in animals spanning the full size range of terrestrial mammals.

Several studies have investigated the compensatory mechanisms that animals could be using to overcome the drawbacks caused by sensorimotor delays. The models in this paper use the simplest method to deal with time delays in a feedback system, which is to reduce the controller gains— this slows down the system’s performance but prevents instability. Weiland et al. (1986) artificially introduced time delays into the reflex loops that control the femur-tibia joint in stick insects, and showed that increasing time delays caused instability in the form of tremors [56]. Tuthill et al. (2018, 2021) proposed that the supraspinal motor centers act to inhibit activity—reduce the controller gains—in reflex circuits in order to keep them stable despite the sensorimotor delays, because severing supraspinal inputs results in tremors in animals [61,62]. A more computationally intensive method to compensate for delays is to use neural prediction—the nervous system could implement internal models that can estimate the present state of the body from time delayed sensory feedback. Miall et al. (1993) suggested that one of the functions of the cerebellum is to act as a Smith predictor, an internal model specifically designed to compensate for time delays in feedback control systems [63]. Rasman et al. (2021, 2023) used a robotic balance simulator to artificially introduce time delays into the sensory feedback signals involved in the neural control of upright standing balance [64,65]. They showed that with training, the internal models involved in upright standing can learn to compensate for even 400 ms of additional time delay, while the human body inherently has only 100–60ms of sensorimotor delays. The nervous system likely encodes several compensatory strategies for sensorimotor delays with varying levels of complexity, with simple and fast strategies mediated through shorter reflex pathways, and slower and more complex strategies mediated through higher (brain stem, cortical, cerebellar) pathways [66].

While we found that sensorimotor delays are detrimental to the control of fast perturbation responses, other studies have pointed out that the nervous system could use time delays to its benefit in certain control situations. Studies have shown that time delayed positive feedback can help stabilize reflexive control pathways, and aid in rhythmic locomotion [48,50,67]. Nishikawa et al. (2007) show that time delays increase gains at the resonant frequency of a control system, which could assist in the control of rhythmic movements [55]. Besides time delays, the nervous system also must deal with other drawbacks such as noise, muscle force capacity limits, limited sensory resolution and sensory dead zones. Milton and colleagues suggest that the nervous system is able to use the interaction between these various drawbacks to simplify neural control [25,68–71]. However, during our simulations, delays did not provide any benefits. For example, if an animal trips during early swing phase while running at high speed, it would be best to react immediately, fully activating its muscles to rapidly and accurately place its foot forward and regain stability. Irrespective of the control strategy, the animal would lose the time required to sense the perturbation, compute a response strategy and transmit action potentials through the reflex pathways. Under feedback control, the animal is further limited, as it must use low gains in order to ensure a stable response, while the corrective movement might not be quick enough to prevent a fall. Under this scenario, and others like it, delays appear to only negatively affect the control of movement.

One implication of our findings is that the compensatory mechanisms used by animals to overcome the disadvantages caused by long sensorimotor delays may vary with size. Previous studies in small animals such as guinea fowl [8,9] and cockroaches [14,15,46] have shown that smaller animals rely more on feedforward control than feedback control for fast locomotion, and utilize the inherent mechanical properties of their musculoskeletal system (preflexes) to compensate for perturbations [72]. This emphasis on feedforward control over feedback makes sense for lessening the consequences of time delays that we have shown here, but feedforward control has its own challenges because accuracy and speed are susceptible to errors in modeling one’s own body, and the perturbations that are being applied. Garcia et al. (2000) showed that as animal size decreases, the damping ratio of their limbs increases—making small animals overdamped and larger animals underdamped [46]. Overdamping will help prevent overshoot or oscillations when repositioning the body making small animals less sensitive to modeling errors in feedforward control. This is a candidate explanation for why smaller animals emphasize feedforward control, which can produce faster responses than feedback control while overdamping still ensures stable movement. Larger animals would not benefit from this damping—their underdamped bodies would be less tolerant to imperfect feedforward control with even small modeling errors resulting in overshooting and oscillations. Therefore, larger animals may emphasize feedback control to ensure accuracy and stability. Our results here further support different control strategies for small and large animals by showing that uncompensated feedback control is up to four times slower than feedforward control in the smallest animals, but only about two to one and a half times slower in the largest animals.

If larger animals could compensate for sensorimotor delays and reduce feedback response times to be shorter than available movement time, feedback control could become viable for fast perturbation responses. Larger animals do not have highly damped joints, but synaptic delays make up a smaller proportion of overall response time [19]. These relatively shorter synaptic delays might allow larger animals to rely on more computational control strategies that combine feedback control with internal models that compensate for delays [63]. Our results show that as animal size increases, delayed feedback control can use more of the available muscle force capacity, narrowing the performance advantage that feedforward control has over feedback control. Effective feedback control would ensure that the underdamped limbs in larger animals are repositioned accurately without oscillations and overshoot/undershoot errors, and when combined with the computational options available when synaptic delays are relatively short, may be a more viable control strategy for larger animals.

## 5. Supporting information

S1 File. Supplementary material. In the document, we have provided detailed derivations for the equations in the paper, and described secondary analyses that we performed to evaluate our results.

S1. Delays in the feedforward and feedback pathways

S2. Bode plot analysis of a linear feedback control system

S3. Normalized feedback control system with time delays and actuator force capacity—detailed derivations and analyses

S4. Normalized feedback model predictions vs. scaled model simulation results

S5. Components of total applied torque under feedback control

S6. Comparing swing and posture task responses to in-vivo perturbation studies

## Supplementary materials

### S1. Delays in the feedforward and feedback pathways

Reflex loops suffer from several sensorimotor delays (*t*_*SM*_) which are distributed across the feedforward (motor) and feedback (sensory) pathways [1]. Between the sensors (e.g. cutaneous receptors, muscle spindles) and the spinal synapse with the motor neuron, there are sensing delays, nerve conduction delays and synaptic delays—we refer to these delays together as sensory delays (*t*_*SD*_). Between the spinal synapse and muscles, there are nerve conduction delays, neuromuscular junction delays, electromechanical delays and force generation delays—we refer to these delays together as motor delays (*t*_*MD*_). In the feedback control model, we placed all the sensorimotor delays in the feedback (sensory) pathway to simplify the analyses (Fig 1). We could instead have placed the motor delays in the feedforward (motor) pathway, and the sensory delays in the feedback (sensory) pathway. Here, we briefly discuss how this assumption does not affect our estimates of response time, as it doesn’t affect settling times for a step response. The closed loop transfer function of a system *J*(*s*), with a transfer function *G*(*s*) in the feedforward (motor) pathway and *H*(*s*) in the feedback (sensory) pathway is given by 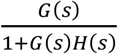 Therefore, the block diagram in Fig S1 has the transfer function:

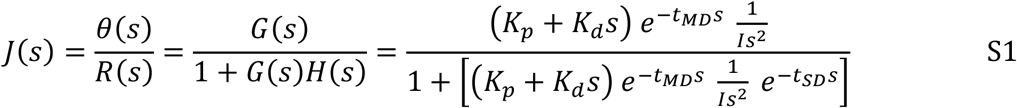

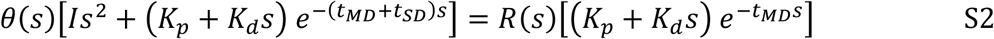

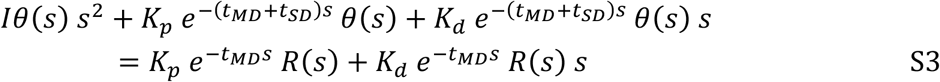

Converting Eqn S3 from the *s* domain to the time domain:

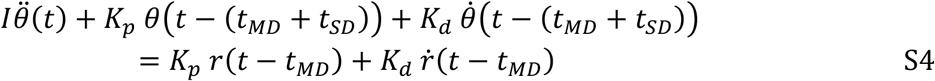

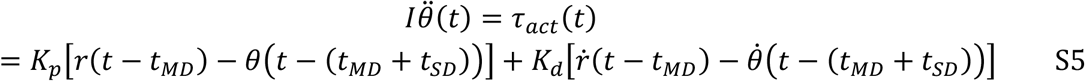

**Fig S1.**
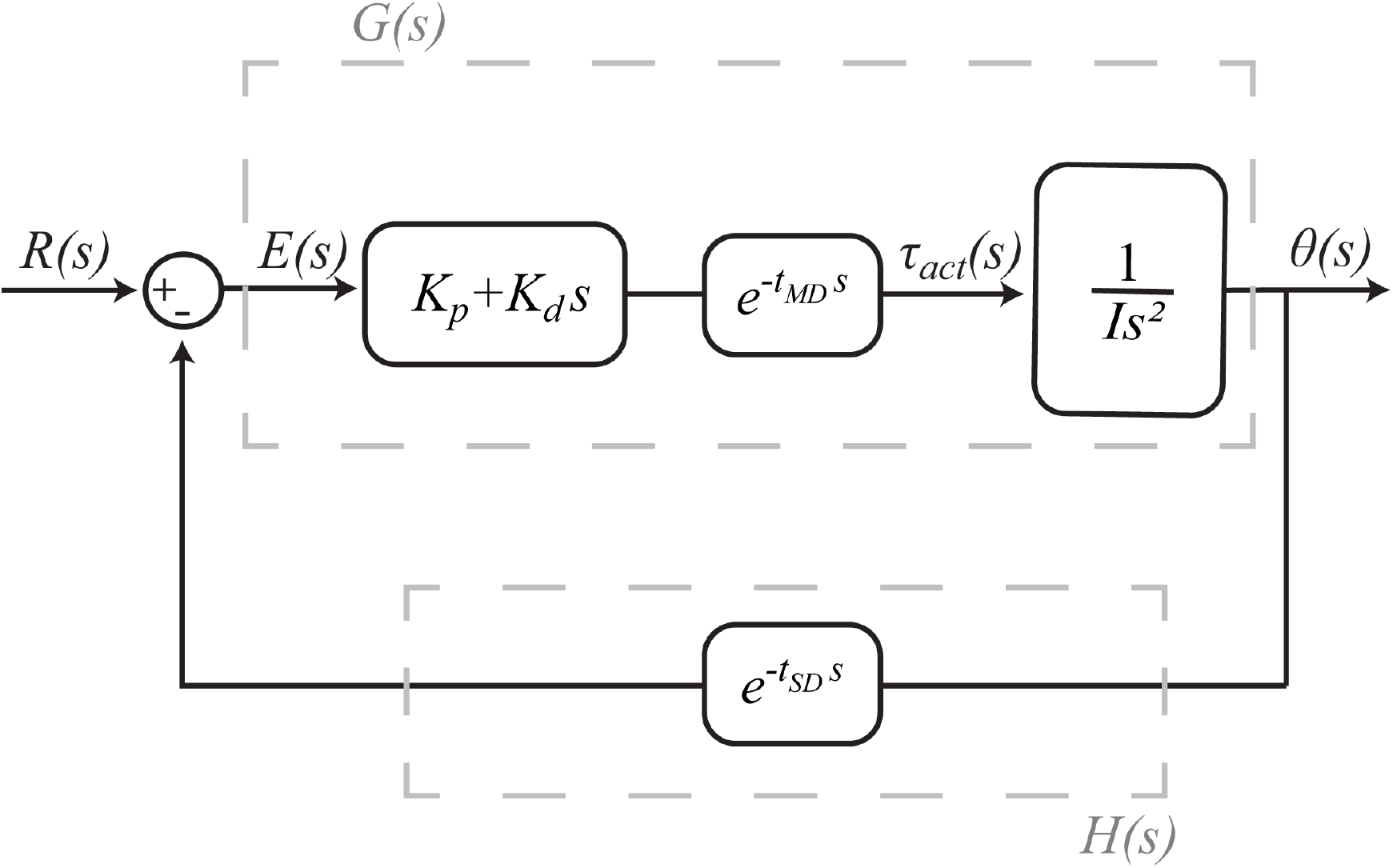
Block diagram with delays in feedforward and feedback pathways. *R*(*s*) is the reference signal, *E*(*s*) is the error signal, *θ*(*s*) is the plant output (angle of the pendulum represented by the double integrator), and *τ*_*act*_(*s*) is the actuating torque. *K*_*p*_ and *K*_*d*_ are the controller gains, and 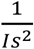 represents the double integrator plant.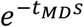 is the transfer function that delays the motor commands due to the time delay *t*_*MD*_, and 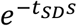 is the equivalent transfer function for the sensory feedback.

The total sensorimotor delay is the sum of the sensory and motor delays:

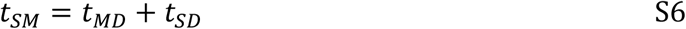

Eqn S5 demonstrates that when there are separate delays in the sensory and motor pathways, the reference signal is delayed only by motor delay (*t*_*MD*_), while sensory feedback is delayed by the total delay (*t*_*SM*_).

However, if we combine all the delays and put them in the feedback (sensory) pathway, we could remove *t*_*MD*_ from Eqn S5 and replace *t*_*SD*_ with *t*_*SM*_ to get:

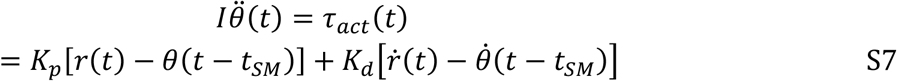

A comparison of Eqns S5 and S7 clarifies how having delays in both the feedforward and feedback pathways will make the tracking of a time varying reference signal more difficult vs. if there were delays only in the feedback pathway. However, since we only consider a step response in our simulations, the reference signal is constant. Therefore, Eqn S5 and S7 are equivalent.

### S2. Bode plot analysis of a linear feedback control system

Here, we revisit a textbook example from linear control systems theory to explain how time delays limit the controller gains that can be used in a stable feedback system. Bode plots are a frequency domain tool for linear feedback systems that determine stability margins; these margins represent the range of parameters that one can use without destabilizing the system. For this analysis, we linearized the scaled models in section 2.1 by ignoring gravity, removing time delays and force capacity limits, and simplifying the plant to a double integrator (a dynamical system representing a single degree-of-freedom rotational system). We derived and analyzed the equations using MATLAB’s symbolic toolbox. Fig S2 shows a block diagram of the linear feedback control system. It represents a Proportional-Derivative (PD) controlled double integrator, and the second-order differential equation of this system is given by:

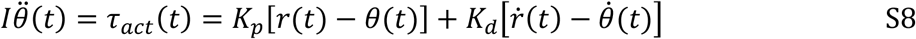

where *I* is the moment of inertia of the plant, and *K*_*p*_ and *K*_*d*_ are the controller gains. Note that *r*(*t*) represents the sinusoidal reference signals used for Bode plots, which is different from *θ*_*r*_ used in the scaled models (the constant target angle for the perturbation response). 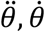 and *θ* are the acceleration, angular velocity and angle of the plant, respectively. The controller produces torques *τ*_*act*_ to reduce the error between the reference signal and the current state, *e*(*t*) = r(t) ™θ(*t*).

We performed a Laplace transform on Eqn S8 to convert it from the time domain to the frequency domain, e.g. *r*(*t*) → R(s), in order to get a transfer function that maps the commanded frequency *R*(*s*) onto the plant output *θ*(*s*). The open loop transfer function of the control system is given by:

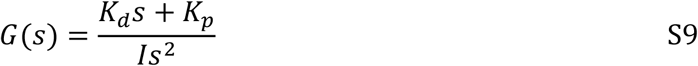

The closed loop transfer function of the control system with unity feedback is given by:

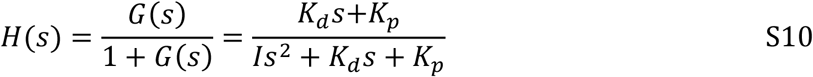

The equation for this closed loop control system is equivalent to that for a mechanical mass-spring-damper system. We derived two more parameters which describe the behavior of the system—the undamped natural frequency 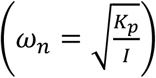 and the damping ratio 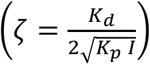 [2–4]. In order to simplify the system, we assumed that the moment of inertia (*I*) is constant, and that the system is critically damped. A critically damped system has a damping ratio of 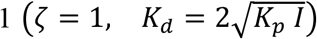, and produces the fastest response without any overshoot [4]. While the scaled models had nine parameters (*I, M, g, L, K*_*p*_, *K*_*d*_, *τ*_*iso*_, *t*_*SM*_, *θ*_*r*_), the linear system initially had three (*I, K*_*p*_, *K*_*d*_), and we have further reduced the system to just a single free parameter (*K*_*p*_).

**Fig S2.**
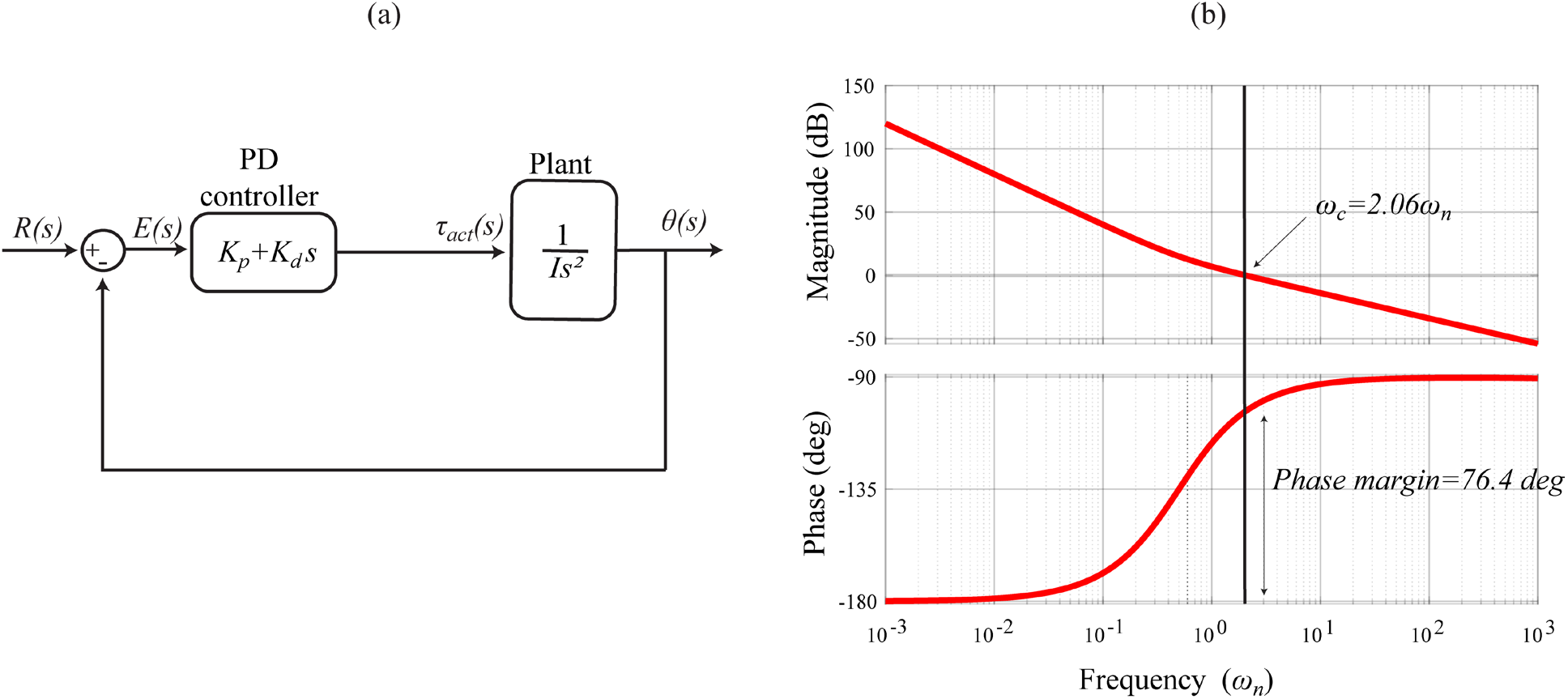
Linear feedback control system and Bode plot. (a) The block diagram of the linear control system, where *R*(*s*) is the reference signal, *E*(*s*) is the error signal, *θ*(*s*) is the plant output (angle of the plant represented by the double integrator), *τ*_*act*_(*s*) the actuating torque, and *s* indicates that these time-dependent variables are represented in the frequency domain. (b) Bode plot of the open loop transfer function (Eqn S9) with the gain plot on top and the phase plot on the bottom. We also show the gain crossover frequency (Eqn S12) and phase margin (Eqn S14) on the plot.

A Bode plot of the open loop transfer function provides the stability margins of the control system (Fig S2b). If the open loop transfer function *G*(*s*) equals -1, the closed loop transfer function in Eqn S10 becomes ∞, signifying instability. This occurs when *G*(*s*) has a magnitude of 0 dB and a phase of -180° for any input frequency in the Bode plot. In this condition, the feedback signal will actively amplify the error and destabilize the feedback control system, instead of attenuating the error and stabilizing it. The phase margin is the distance of the phase line (Fig S2b bottom) from -180°, at the gain crossover frequency (*ω*_*c*_) — the frequency where the gain line crosses the 0 dB line (Fig S2b top). We found the gain crossover frequency by solving for the frequency at which the gain of the open loop transfer function equaled 1 (0 dB).

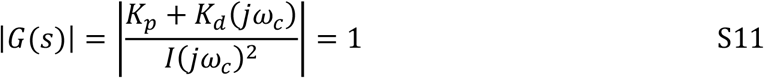

Substituting in the value of *K*_*d*_ for critical damping and solving for *ω*_*c*_ gives:

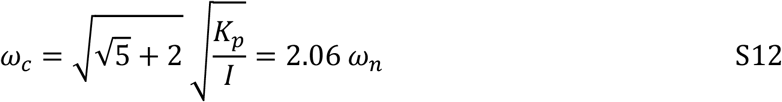

The phase margin is how far the phase of the open loop transfer function at the gain crossover frequency is from 180°:

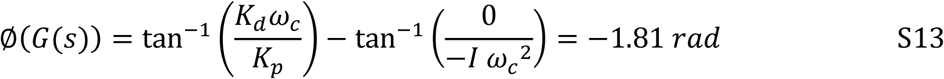

Eqn S13 reduces to a constant value because substituting in Eqn S12 for *ω*_*c*_ and the critical damping equation for *K*_*d*_ caused all variable terms to cancel out.

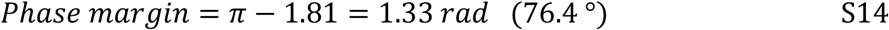

If this feedback system is subjected to a phase shift that exceeds the phase margin, it will become unstable. Time delays in the signal pathways can cause a phase shift, and the time delay that corresponds to the phase margin is called the delay margin. We obtained the delay margin by dividing the phase margin by the gain crossover frequency:

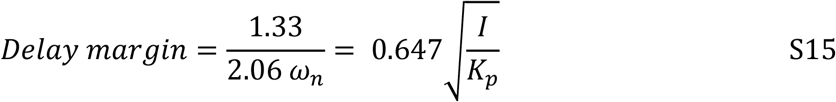

As animals have a fixed sensorimotor time delay within their reflex pathways (*t*_*SM*_), we can invert Eqn S15 to show that the maximum proportional gain 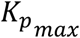 that a control system with a given time delay *t*_*SM*_ can use is:

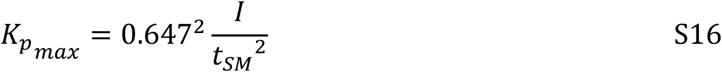

At 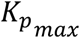, we can also determine *K*_*d*_ and *ω*_*n:*_

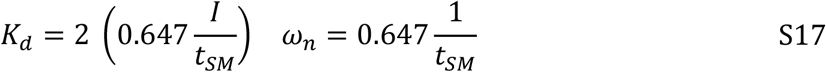

Thus, for a feedback control system with time delay to remain stable, it must limit its controller gains such that:

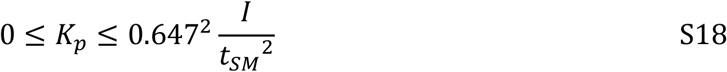

Eqn S17 shows that the time delay will cap the maximum natural frequency of the feedback system. We also find the following associations between the parameters of the feedback system: 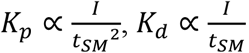 and 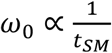.

### S3. Normalized feedback control system with time delays and force capacity limits—detailed derivations and analyses

We developed a normalized feedback control system (one model which can represent animals of all sizes), and studied how response times are affected by time delays and muscle force capacity (saturation) limits (Fig S3). To keep the paper concise, the main manuscript describes an abridged version of this analysis. Here, we have provided the detailed description and complete text of this analysis. The equations of motion of the feedback control system before normalization are:

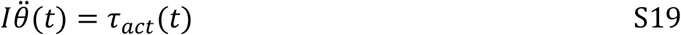

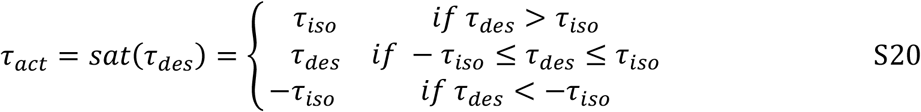

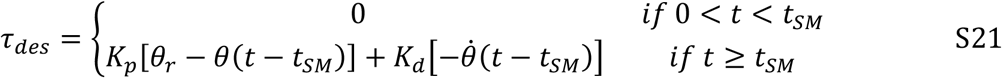

where τ_*des*_ is the desired controller output. *τ*_*act*_ is the actual torque applied to plant, subject to force capacity limits *τ*_*iso*_. We did not consider actuator (muscle) dynamics in our models, and directly applied the controller output as torques to the plant. We also assumed that the controller gets full state information; we did not consider sensory dynamics. Due to time delays *t*_*SM*_, there is an initial deadtime at the start of the simulation where no torque is applied to the plant (Eqn S21). We only consider a constant reference target *θ*_*r*_ in these simulations.

We normalized Eqns S20 and S21 using three constants which also represent characteristic features of the neural control of movement in animals:

*I* is the moment of inertia of the double integrator plant, and also represents the moment of inertia of the body segments being moved under neural control by the animal.

*θ*_*r*_ is the reference target for feedback control, and also represents the size of the movement being commanded under neural control in the swing task. For the posture task, the movement size is represented by the initial perturbation velocity 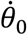.

*t*_*SM*_ is the time delay in the feedback control system, and also represents the sensorimotor delays in the reflex pathways during neural control of movement.

**Fig S3.**
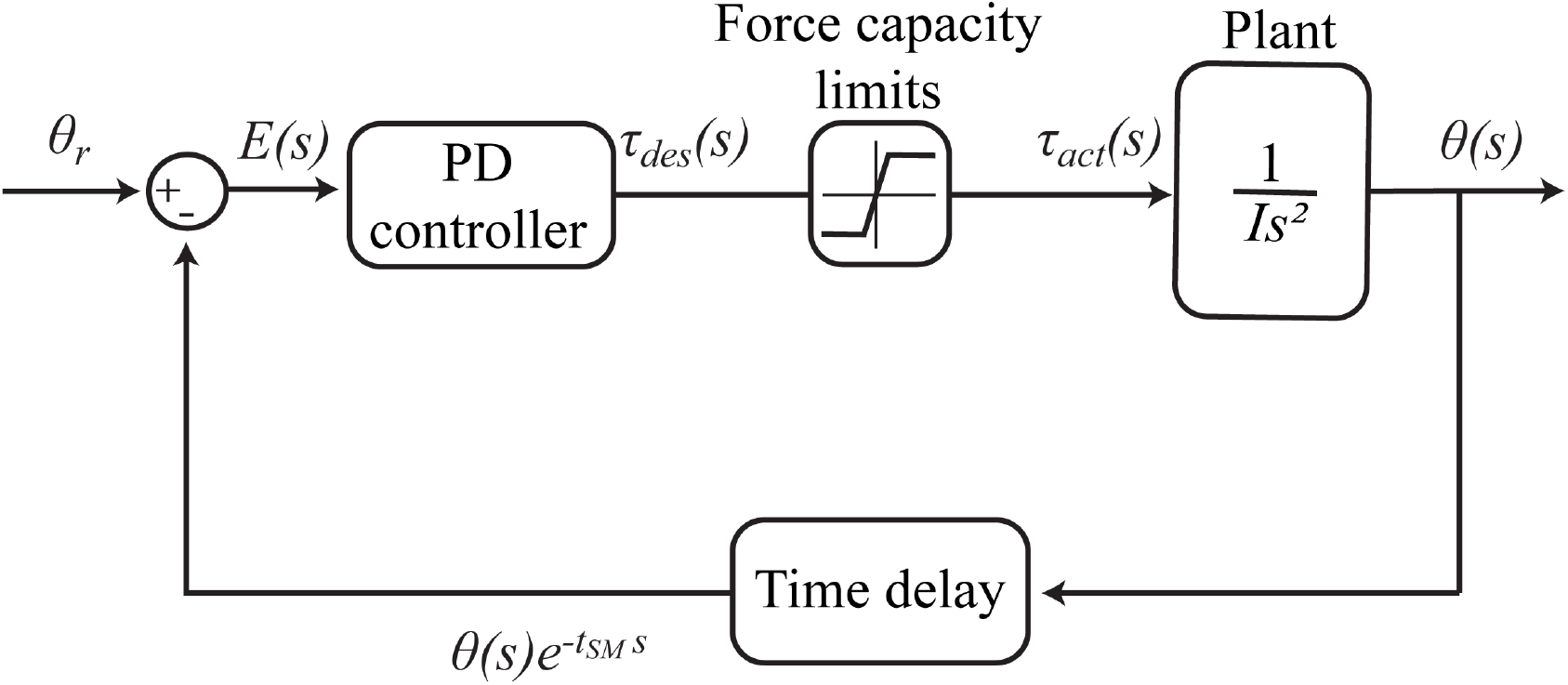
Feedback control system with time delays and force capacity limits. *θ*_*r*_ is the reference target, *E*(*s*) is the error signal, *θ*(*s*) is the plant output (angle of the pendulum represented by the double integrator), *τ*_*des*_ is the controller output torque, and *τ*_*act*_ the actuating torque subject to force capacity limits. 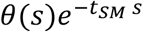 is the time delayed feedback signal.

The terms in Eqn S21 have dimensions of torque (M^1^L^2^T^-2^), where M represents mass, L represents length and T represents time dimensions. Therefore, to derive the normalized model, we divided each term in Eqns S20 and S21 by 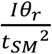.

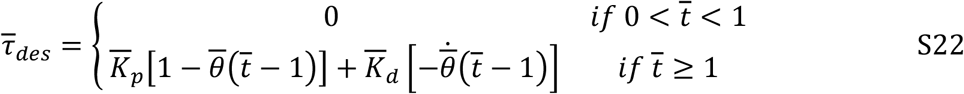

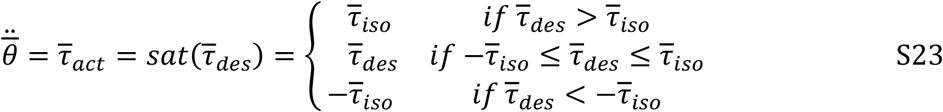

The normalized model is described by Eqns S22 and S23. Below, we describe how to normalize each of the parameters of the model.

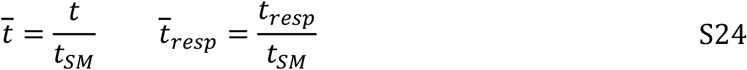

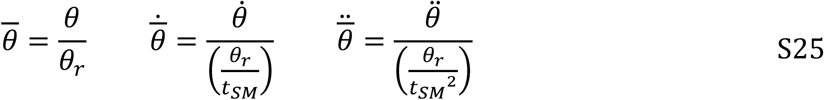

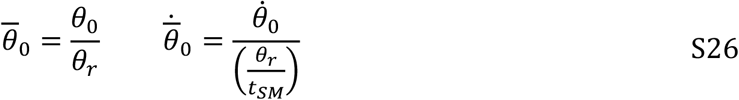

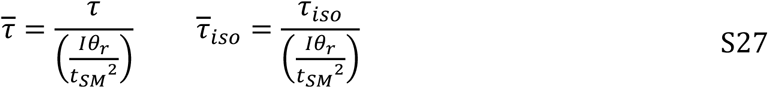

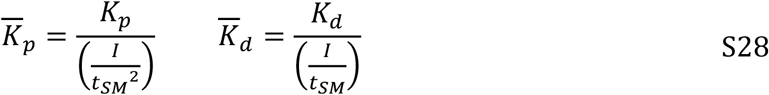

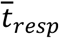 is the normalized response time of the model, determined using numerical simulations and optimization.

#### Normalized feedback control system—numerical simulations

Using numerical simulations, we evaluated how changing the perturbation task, controller gains, time delays and force capacity limits affected the behavior of the normalized feedback control system. First, we did not set force capacity limits, used a fixed time delay 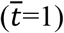, and performed a brute force search to determine how controller gains affect settling time and overshoot. Next, we kept time delays constant 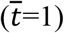, and varied the force capacity limits to understand how this affects response times 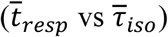. For each force capacity limit, we optimized the controller gains to find the fastest response time (fastest settling time with 2% thresholds and without any overshoot).

#### Swing task results

A brute force search through a range of controller gains revealed the settling time and overshoot landscapes for the swing task. We calculated both settling time and overshoot on the angle curve for the swing task (Fig S4a bottom panel). The settling time landscape is very rough and jagged with several local minima (Fig S4b). The overshoot landscape has a flat area with zero overshoot, followed by a region of rapid increase (Fig S4c). The black dot depicts the fastest settling time achieved while allowing overshoot. The red dot depicts the fastest settling time without overshoot; we used this result to determine response time. Controller gains of 0.1617 for 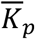 and 0.6343 for 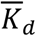 produced the fastest response time 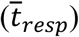 of 7.09 in normalized time units (Fig S4a).

Analysis of the relationship between force capacity limits and response time revealed three distinct regions (Fig S5). The fastest response without considering force capacity limits produced a peak torque of 0.1617, which equals 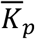. Fig S5 shows how response time, overshoot and controller gains changed when we varied force capacity limits from 0.25 to 0.001.

- High force capacity limits region: For 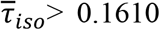, the peak torque commanded by the controller did not reach the force capacity limits, and the response time and controller gains did not change.
- Middle region: For 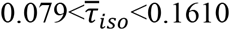, the force capacity limits clipped the positive region of the torque curve, and response time increased gradually with lower force capacity limits. However, the optimal control gains still remained the same.
- Low region: 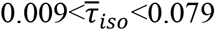, the force capacity limits clipped both the positive and negative regions of the torque curve, and response time increased exponentially with lower force capacity limits. In this region, the controller gains also increased with lower force capacity limits.

**Fig S4.**
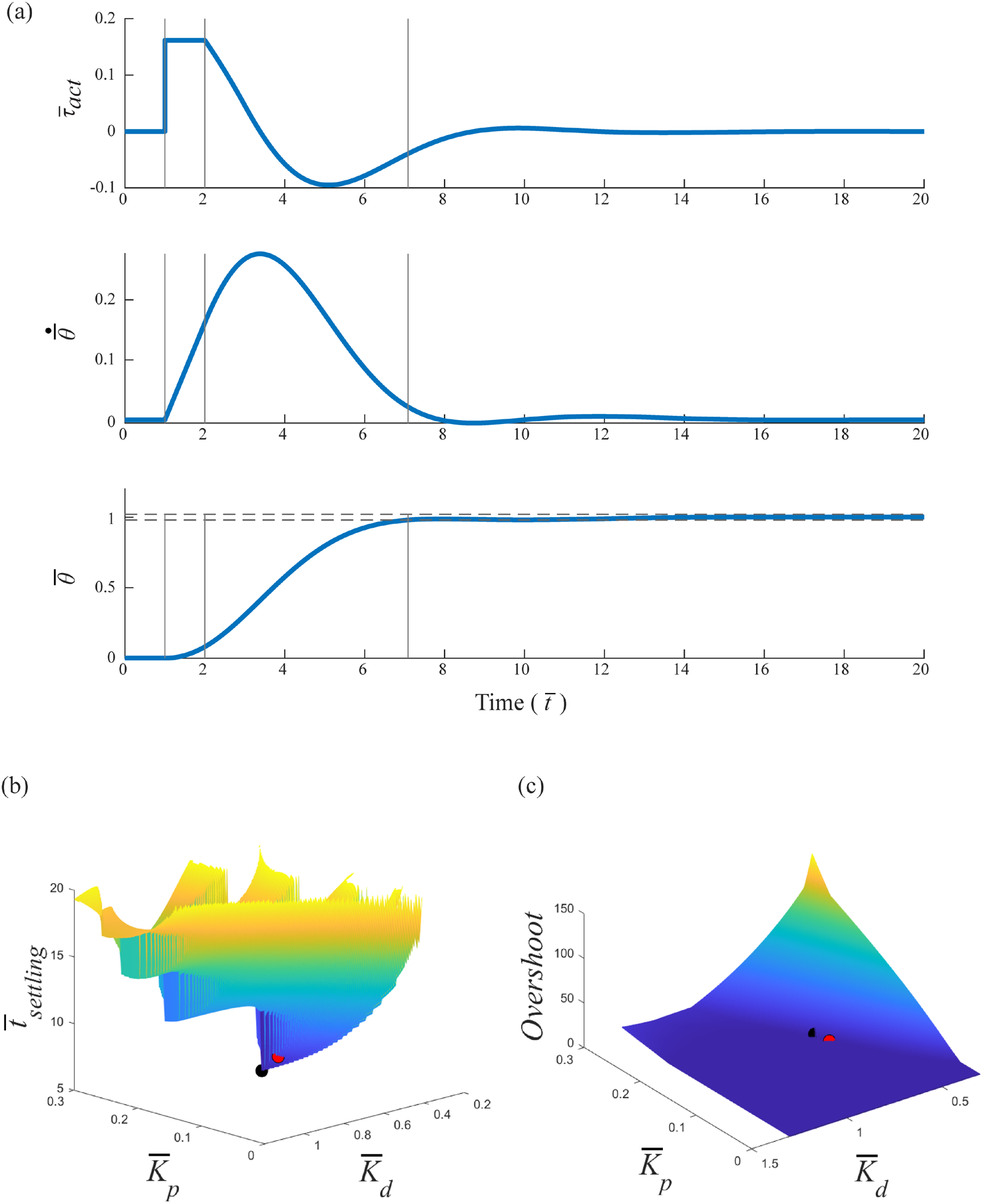
Normalized swing task—brute force search. (a) Normalized torque 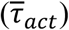, angular velocity 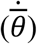 and angle 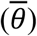 profiles in the swing task for the fastest response. The three grey vertical lines depict one time delay period, two time delay periods and the settling time. The two grey horizontal dashed lines on the angle graph depict the 2% settling time thresholds. (b & c) The settling time and overshoot landscapes for the normalized swing task model determined through a brute force search for a range of controller gains. The black dot depicts the fastest settling time when overshoot is allowed (This response had 1.76% overshoot). The red dot depicts the result we use for response time—the fastest settling time without overshoot.

**Fig S5.**
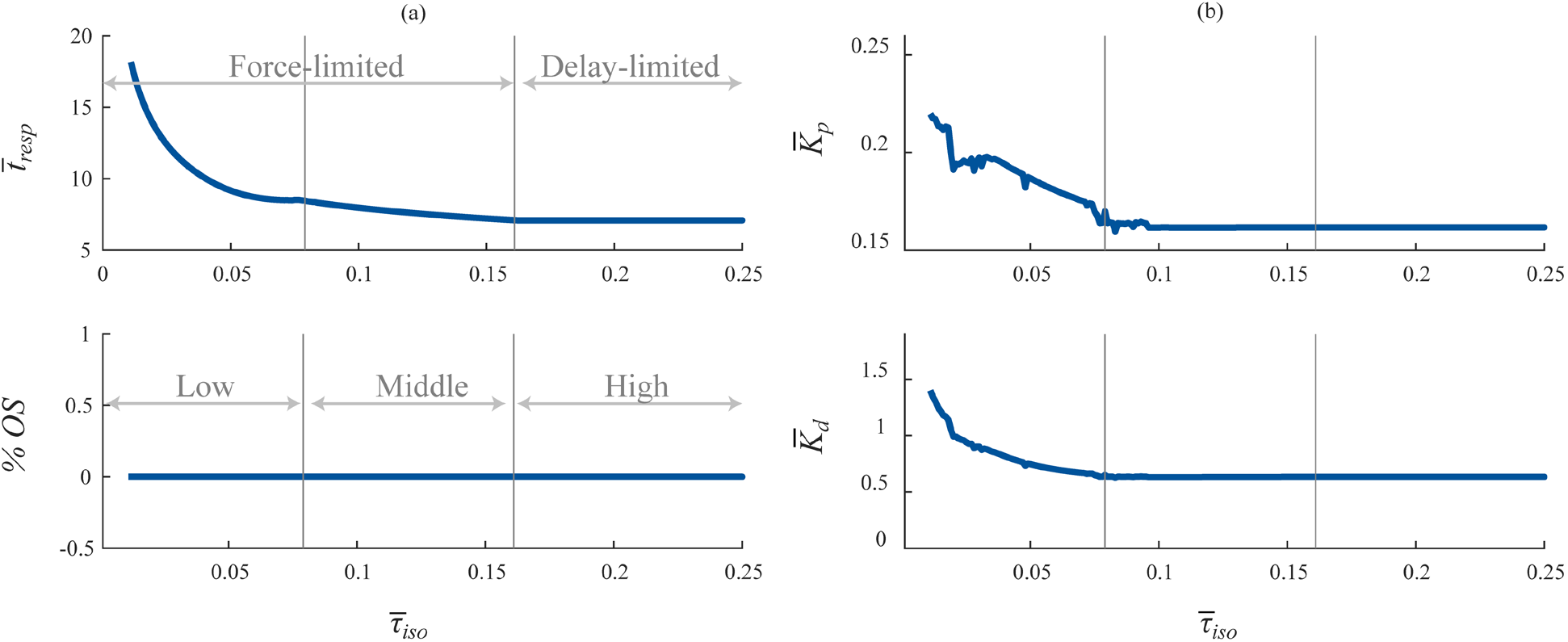
(Fig 2). Normalized swing task—force capacity limits vs. response time. (a) Changes in response time and overshoot when force capacity limits are lowered from 0.25 to There are three distinct regions in the response time profile, shown divided by the grey vertical lines. (b) Changes in the optimal controller gains that produce the fastest response times without overshoot for a range of force capacity limits. Note that there is some variability in the optimal controller gains in the middle and low regions due to multiple local minima very close to the global minimum, but they do not affect the response time significantly.

To evaluate whether the normalized model can predict response times in the more detailed scaled model simulations, we used curve fitting to estimate the relationship between normalized force capacity and normalized response time 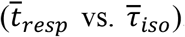. We found that the double exponential function gave the best fit (lowest root mean square error) for the middle and low force capacity limit regions in the swing task. The swing task 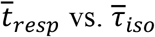 relationship can be described by the following function:

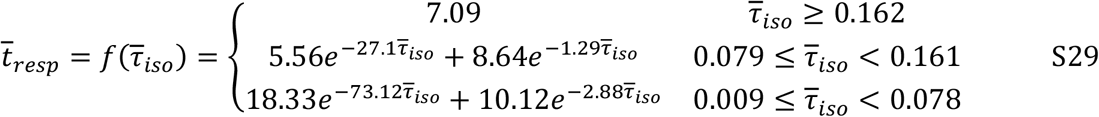

This analysis shows that for feedback control systems with both time delays and force capacity limits, there are two ranges in the response time vs. force capacity limits graph: a force-limited range and a delay-limited range. At the extremes, without force capacity limits or delays, infinitely high gains can produce instant response times. If we reduced the force capacity limits to 0, or if the time delays were infinitely long, we would have infinite response times. The delay-limited range matches the high force capacity limits region 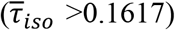, where the response time is limited purely by time delays. The force-limited range consists of the middle and low force capacity limits regions 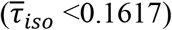, where the force capacity limits also begin to limit response time. Extrapolating to animals, this analysis indicates that for an animal to respond quickly, it requires both strong muscles and short sensorimotor delays. Deficiencies in either factor will slow the animal’s ability to respond quickly. Whether an animal is delay-limited or force-limited would depend on the relative magnitudes of factors that affect the perturbation response, such as the moment of inertia of the body segments being moved, the sensorimotor delays, the size of the perturbation response movement, and the muscle force capacity.

#### Posture task results

For the normalized posture task model, the brute force search results were similar to the swing task, while the relationship between response time and force capacity limits had several differences. For the posture task, we calculated settling time on the angular velocity curve, and overshoot on the angle curve. We did this to ensure that the settling time thresholds scaled with perturbation size, while not changing with controller gains. The settling time landscape was again very rugged. The red dot in Fig S6b depicts the controller gains that produced the fastest settling time; this set of controller gains also caused no overshoot. Without setting force capacity limits, controller gains of 0.1560 for 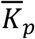 and 0.6530 for 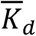 produced the fastest normalized response time 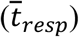 of 7.38 in normalized time units (Fig S6a).

Analysis of the relationship between force capacity limits and response time again revealed three distinct regions. The fastest response without considering force capacity limits produced a peak torque of 0.80. Fig S7 shows how response time, overshoot and controller gains changed when we varied force capacity limits from 1.2 to 0.001.

- High force capacity limits region: For 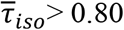, the peak torque 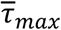 did not reach the force capacity limits, and the response time and controller gains did not change.
- Middle region: For 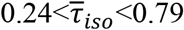, the force capacity limits clipped the positive region of the torque curve, and response time increased gradually with lower force capacity limits. Unlike the swing task where the controller gains did not change in the middle region, the controller gains increased gradually with lower force capacity limits.
- Low region: 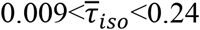, the force capacity limits clipped both the positive and negative regions of the torque curve. Unlike the swing task, the response time continued to increase at the same rate as in the middle region. The controller gains initially show a shallow dip before increasing rapidly for lower force capacity limits.

**Fig S6.**
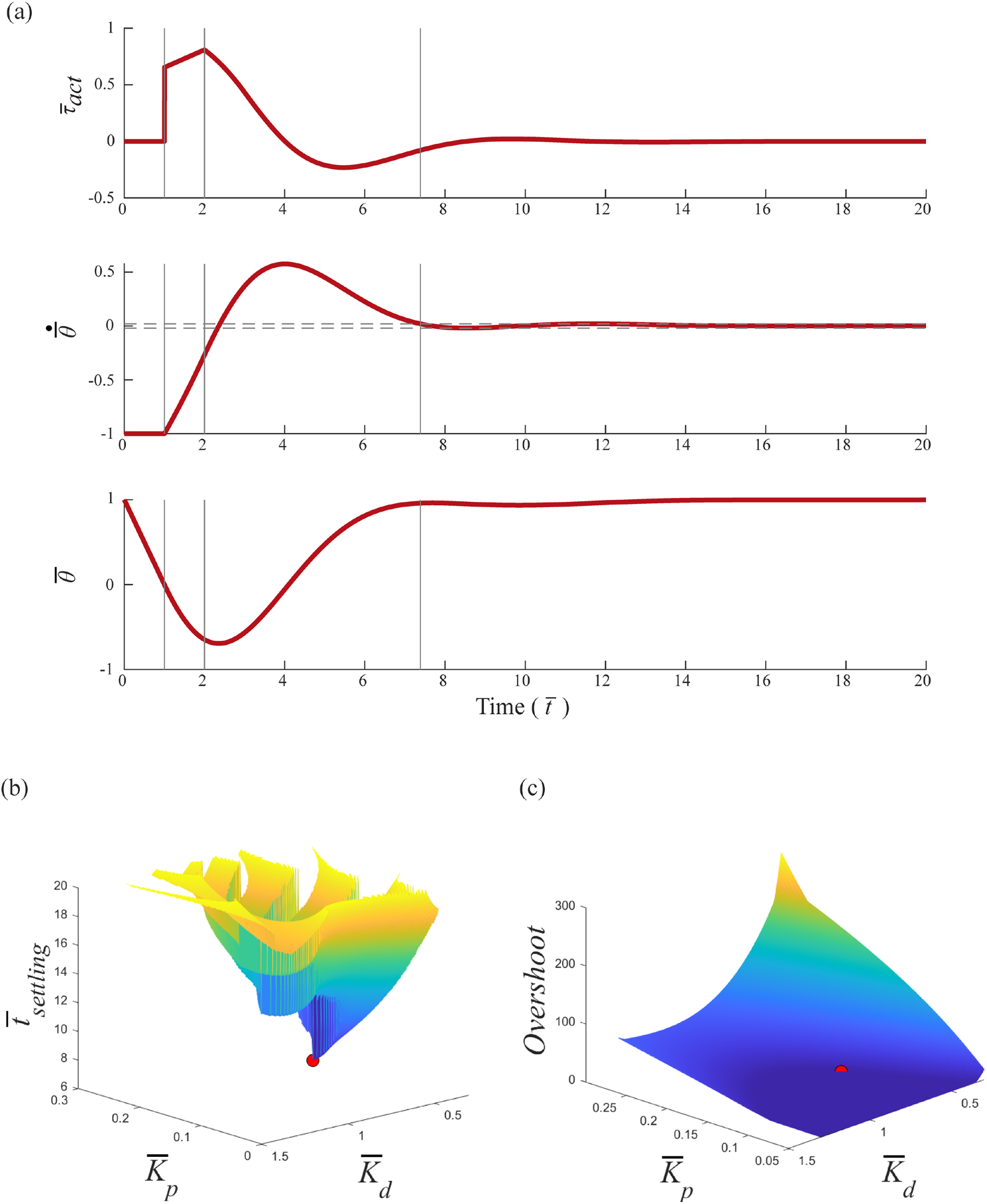
Normalized posture task—brute force search. (a) Normalized torque 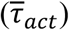, angular velocity 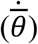 and angle 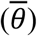 profiles in the posture task for the fastest response. The three grey vertical lines depict one time delay period, two time delay periods and the settling time. The two grey horizontal dashed lines on the angular velocity graph depict the 2% settling time thresholds. (b & c) The settling time and overshoot landscapes for the normalized posture task model determined through a brute force search for a range of controller gains. The red dot depicts the result we use for response time—the fastest settling time. The value does not change if overshoot is allowed or constrained.

**Fig S7.**
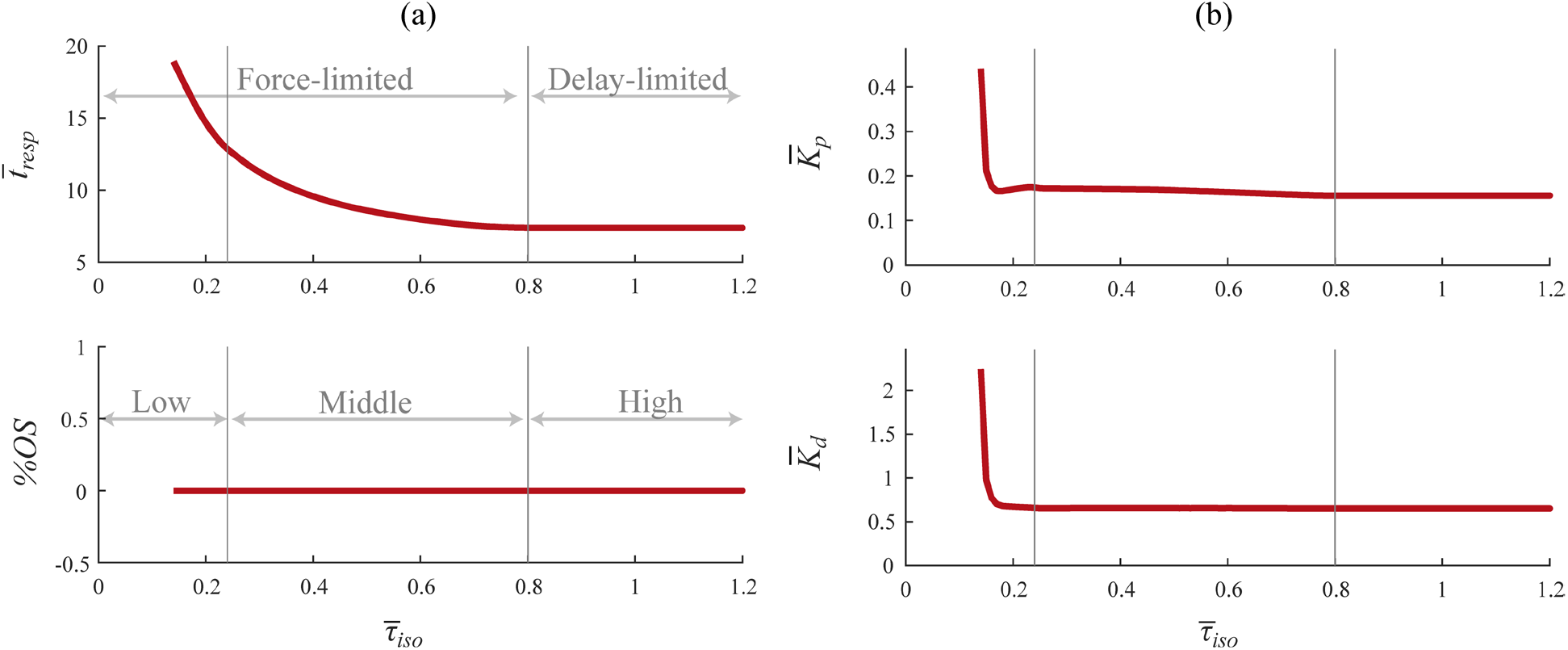
Normalized posture task—force capacity limits vs. response time. (a) Changes in response time and overshoot when force capacity limits are lowered from 1.2 to There are three distinct regions in the response time profile, shown divided by the grey vertical lines. (b) Changes in the optimal controller gains that produce the fastest response times without overshoot for a range of force capacity limits.

We again tried fitting various functions to the settling time curve in the middle and lower regions. We found that a power law with intercept (power2) function produced the lowest RMSE values. The posture task 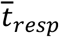 vs .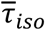 relationship can be described by the following function:

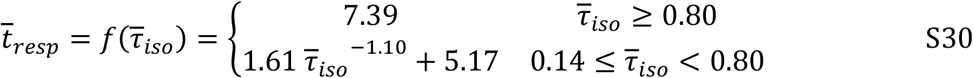

### S4. Normalized feedback model predictions vs. scaled model simulation results

Predictions for response time based on the relationship between normalized force capacity limits and response time 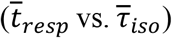 from the normalized model (Section S3) compared well to results from the more detailed scaled model simulations (Section 3.2), despite their differences. The normalized models did not incorporate gravitational torques, while the scaled models considered them. Additionally, the swing task scaled simulations incorporated a steady state torque to counter gravitational torque at the target state. Feedback control response times in the scaled simulations for the swing task scaled as 199 *M*^0.21^ ms. The normalized equations (Eqn S29) predicted response time from the scaled simulations with an average accuracy of -10.5% (range: -8%, -14%) (Fig S8 left). Feedback control response times in the scaled simulations for the posture task scaled as 239 *M*^0.22^ ms. The normalized equations (Eqn S30) predicted response time from the scaled models with an average accuracy of 5% (range: 2%, 13%) (Fig S8 right).

**Fig S8.**
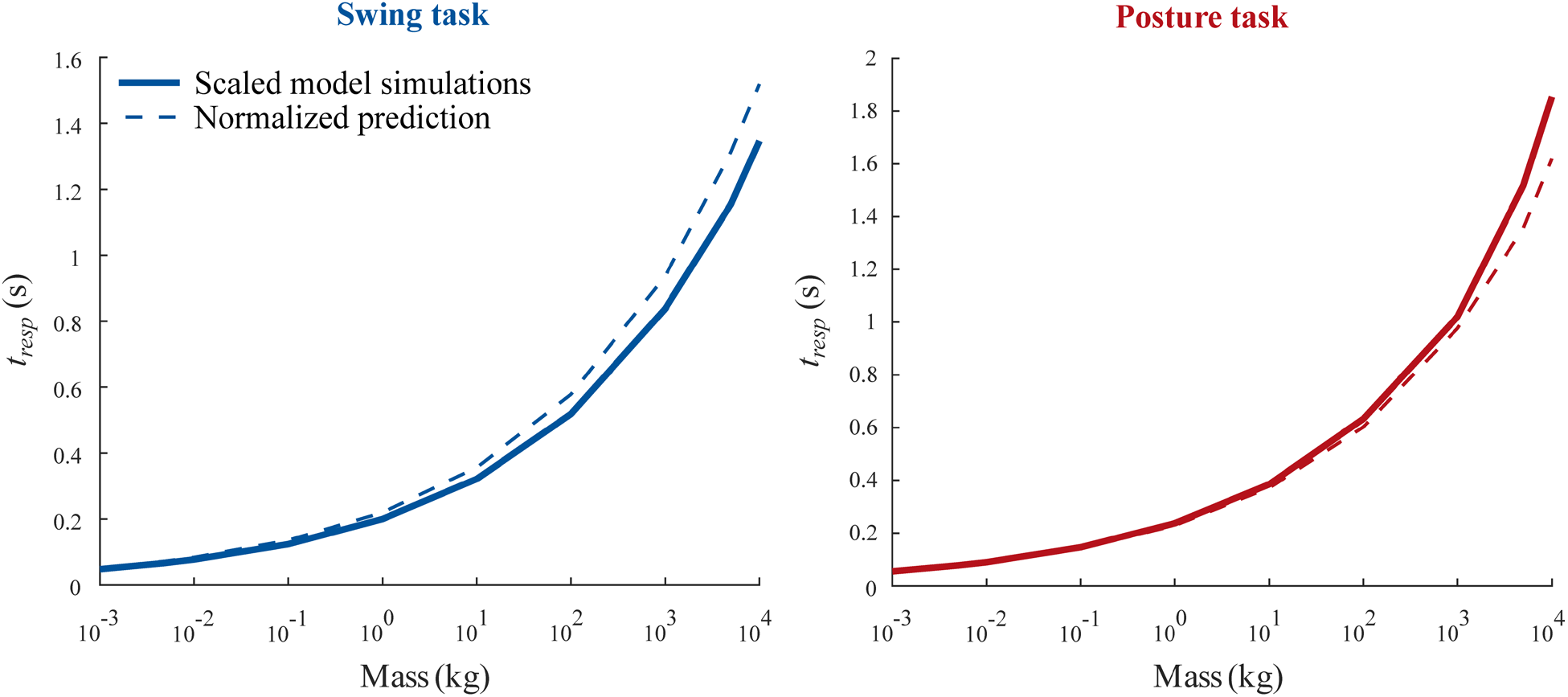
Feedback response time—normalized model predictions vs. scaled simulation results. Normalized prediction for the swing task (left) and posture task (right) in dotted lines. Scaled simulation results in solid lines.

### S5. Components of total applied torque under feedback control

We calculated the total torque experienced by the plant as the sum of the muscle torque (generated by the feedback controller), and gravitational torque. The muscle torque composed of proportional and derivative components in the posture task, and an additional steady state component in the swing task. We capped the muscle torques at a force capacity limit, computed as the maximum isometric torque that can be produced by the relevant muscles. Applied torques exceeded force capacity limits only for animals heavier than one ton in the posture task. During an initial delay period equal to the sensorimotor delay, the controller did not apply muscle torques and the plant moved purely under gravitational torque. We considered this the time required for the animal to sense the perturbation, compute the motor commands, and transmit the signals to the muscles. After this deadtime, the controller turned on the muscle torques, computed based on time delayed state feedback. Fig S9 depicts the contribution of each component to the applied torque for a 1 kg animal in the swing task and posture task.

**Fig S9.**
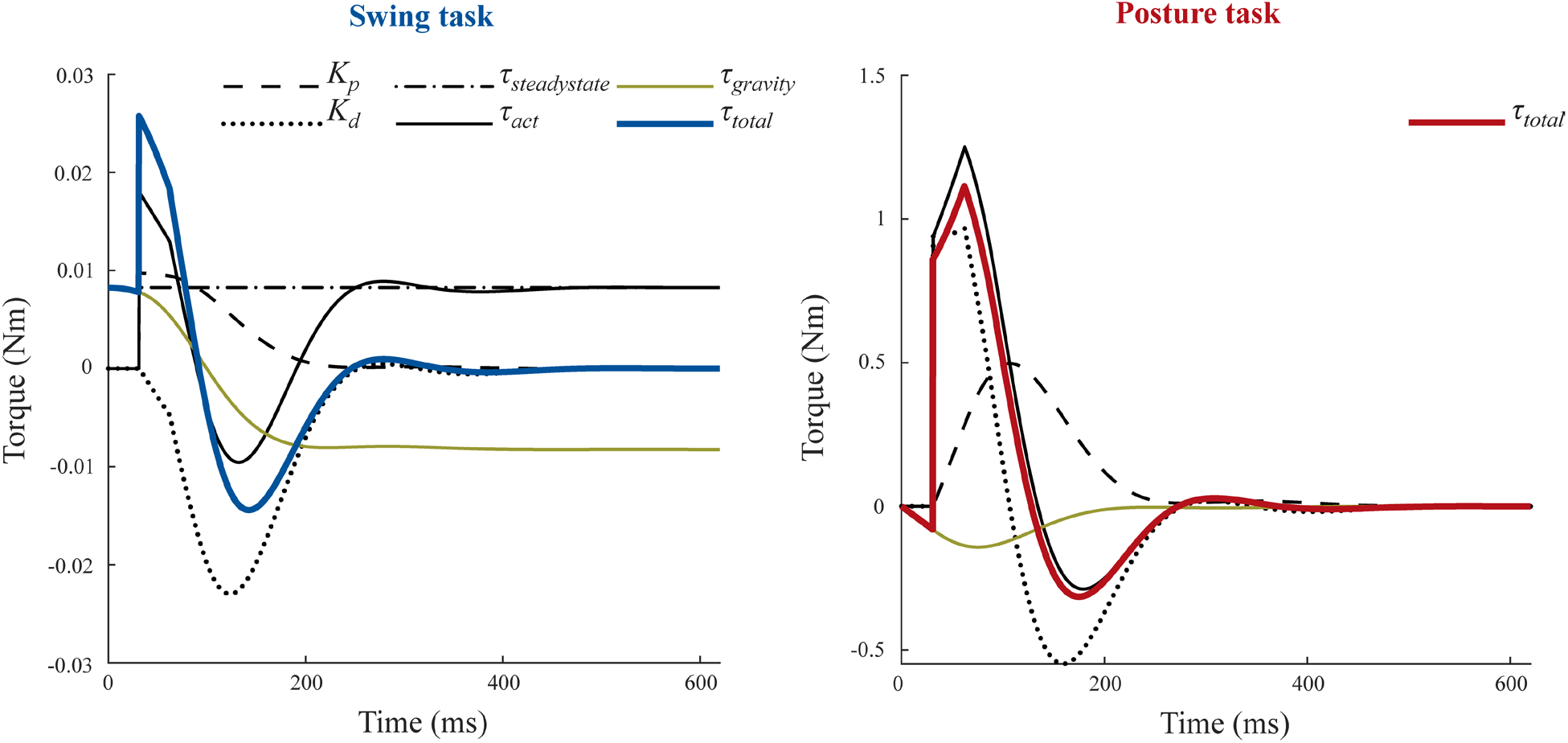
Components of total torque. For the swing task on the left, the controller torque consisted of a proportional component (*K*_*p*_— black dashed line), a derivative component (*K*_*d*_—black dotted line), and a steady state component (*τ*_*steadystate*_—black dash dot line). These three components made up the muscle torque (*τ*_*act*_— solid black line). The total torque (*τ*_*total*_—thick blue line) is the sum of *τ*_*act*_ and gravitational torque (*τ*_*gravity*_—yellow green line). For the posture task on the right, the total torque is shown in red. The posture task does not have a steady state torque component. The values shown here are for a one kg animal in the swing task and posture task.

### S6. Comparing swing and posture task responses to in-vivo perturbation studies

We compared the kinematic profiles from our simulations to in-vivo perturbation studies, and found that they were qualitatively similar, despite the limitations of our computational models. In-vivo perturbation studies on animals do not elicit the fastest possible responses in order to prevent falling and injury. Our simulations model the fastest perturbation responses controlled purely through monosynaptic reflex pathways, and consider feedforward and feedback control separately. Perturbation responses from in-vivo studies are difficult to separate into purely feedforward or feedback strategies, and often involve both the reflexive and supra-spinal motor pathways. Eng et al. studied stumble corrective responses in humans, and reported that an approximately 40° swing leg repositioning response to an early swing phase trip took about 500 ms [5]. According to the swing task simulations for a 70 kg human performing a 30° swing leg repositioning, the fastest possible responses would take 172 ms under feedforward control and 487 ms under feedback control.

We also compared the posture task simulations to three studies that reported on human and cat postural responses to support-surface translations [6–8]. Horak and Nashner (1986) reported that for a 0.05 dimensionless velocity perturbation, their human subjects took about 600 ms to regain posture, and suffered a maximum lean of 3° [6]. Welch and Ting (2008) reported that for a 0.09 dimensionless velocity perturbation, their human subjects took about 1000 ms to regain posture, and suffered a maximum lean of 5° [7]. The posture task simulations predict that a 70 kg human subjected to a 0.21 dimensionless velocity perturbation would take 310 ms under feedforward control and 611 ms under feedback control to recover balance, and force them to lean to a maximum angle of about 5°. Ting and Macpherson (2004) reported that cats subjected to a 0.09 dimensionless velocity postural perturbation take about 1000 ms to regain posture, and exhibit a 5° lean [8]. The posture task simulations predict that for a 4 kg cat subjected to a 0.21 dimensionless velocity perturbation, recovering balance would take 140 ms under feedforward control and 326 ms under feedback control, and force it to lean to a maximum angle of about 3.5°. Fig S10 shows the angle vs. time profiles for different sized animals in the posture task under feedforward and feedback control. These profiles match the behavior seen in studies on posture correction in quadrupeds. The maximum lean of the posture task model (about 7° for a 0.21 dimensionless velocity perturbation) would not cause the center of mass to move outside the base of support in quadrupeds. However, we have not considered similar effects in bipeds. Bipeds would be forced to use a stepping strategy, instead of a hip or ankle strategy, if large perturbations cause the center of mass to move beyond the base of support.

**Fig S10.**
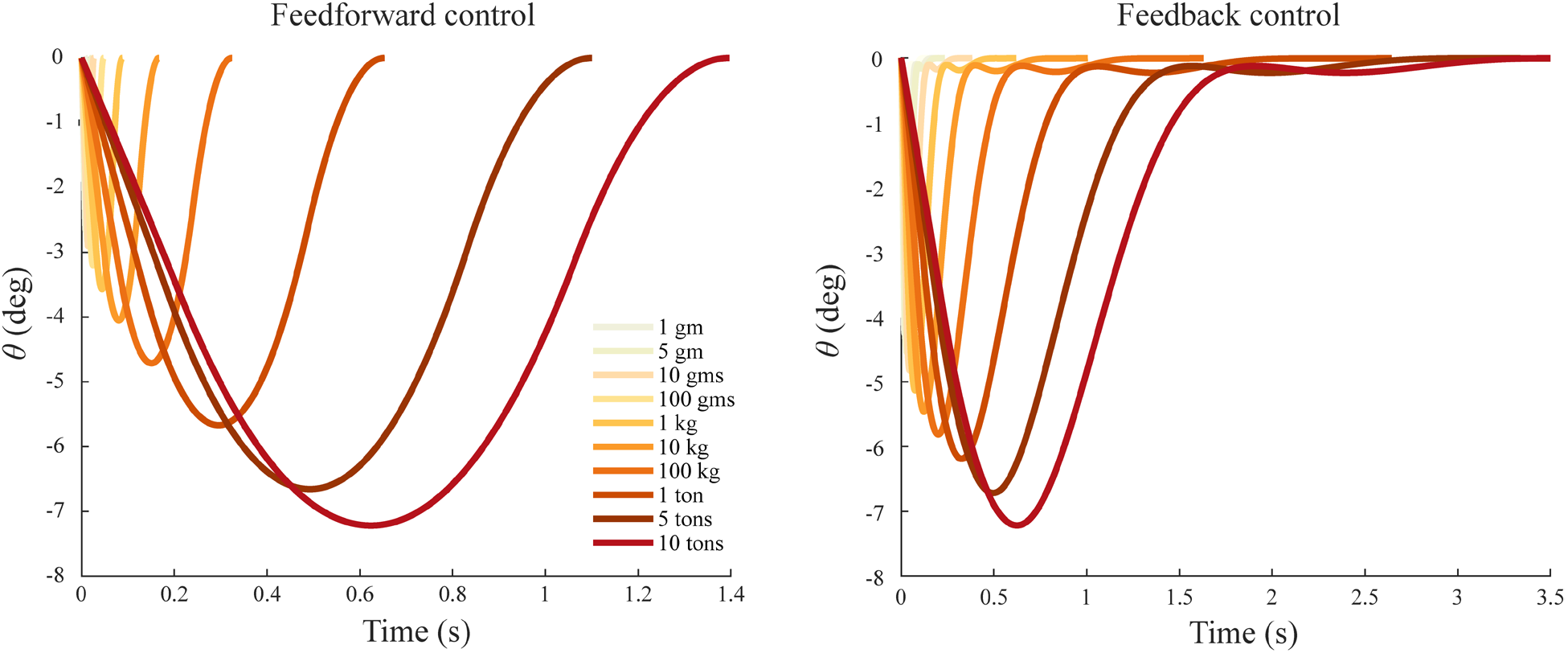
Angle profiles for the posture task under feedforward and feedback control. Angle vs. time profiles from the posture task simulations for animal sizes ranging from 1 gram to 10 tons under feedforward control (left) and feedback control (right), for a 0.21 dimensionless velocity perturbation.

## Notes

### Competing Interest Statement

The authors have declared no competing interest.

